# What we miss shapes what we protect: sea turtles as a case study of grey literature and language bias in an evidence synthesis

**DOI:** 10.64898/2026.07.17.739284

**Authors:** Harris W.K. Heng, Kristine C.V. Buenafe, Jaime Restrepo, Dina Nisthar, Tatsuya Amano, Nicolas Pilcher, The Cuong Chu, Nguyen Trong Duc, Rose Ellis Arnett, Chia-Ling Fong, Sekar M.C. Herandarudewi, Daphne Z. Hoh, Seh Ling Long, Connie Ka Yan Ng, Janmanee Panyawai, Daniel C. Dunn

## Abstract

Evidence-based approaches are increasingly advocated and applied in conservation to provide reliable recommendations for management and policymaking. Although substantial efforts have been undertaken to generate ecological knowledge across languages and publication types, many syntheses and meta-analyses fail to include relevant literature sources; drawing into question their reliability. To assess the extent of biases resulting from ignoring grey literature and non-English publications, we reviewed and synthesised movement and migratory connectivity data for green turtles across Southeast Asia from multiple literature types and eight languages. We identified substantial losses in connectivity information, with at least half of all locations losing more than 88% of their connections, and shifts in demographic representation and sampling techniques when grey literature was excluded from the evidence base. The exclusion of non-English literature had a comparatively smaller impact. The regional connectivity network generated through this study highlights the need for holistic reviews to develop evidence bases to underpin transboundary management of migratory species. Ignoring grey literature and, potentially, regional languages in the development may lead to misallocation of limited management resources. Holistic reviews can further support the establishment of Important Areas for migratory species and inform how we meet and report on global biodiversity commitments.

## 1 Introduction

Evidence-based conservation builds on the synthesis of empirical information produced by individual studies to address contemporary environmental issues. To develop the best solutions, managers and policymakers require a comprehensive understanding of the research field, as well as the extent of current knowledge, and the gaps that may hinder the use of the knowledge in real-world applications (Pullin & Knight, 2012). Ecological knowledge has been continuously and increasingly produced worldwide through scientific endeavours, traditional or indigenous community practices, and modern citizen-science programmes (Fraisl et al., 2020; Hudson et al., 2017; McLean et al., 2023; Reyes-García et al., 2024). However, several barriers exist that prevent decision-makers from applying evidence-based approaches to the conservation and sustainable use of biodiversity (Cook et al., 2012; Dicks et al., 2014a, b), such as limited access to empirical data and gaps between the evidence available and the evidence needed for informed decision-making. Despite centuries of effort to generate ecological knowledge, these barriers continue to hinder its application, contributing to ongoing population declines and species extinctions (Ceballos et al., 2017; IPBES, 2019).

Ecological processes, functions, and species behaviour can vary substantially from one geographical location to another (for examples, see Hill et al., 2025; Mumby et al., 2004). Synthesising evidence across geographic and thus cultural and economic spectrums, and understanding the potential transferability of that evidence, requires gathering information from different types of publications (Haddaway & Bayliss, 2015; Haddaway et al., 2020; Balvanera et al., 2022), in different languages (Konno et al., 2020; Amano et al., 2021; Nuñez et al., 2021; Berdejo-Espinola & Amano, 2025), across many types of institutions with widely varying degrees of accessibility (Driscoll et al., 2021; Nuñez et al., 2021; Balvanera et al., 2022). Thus, the generation of synthetic knowledge via the collation of evidence from multiple sources has to be curated with caution to properly account for biases (Hannah et al., 2025).

Recent studies have shown that conclusions of evidence synthesis can be biased when certain types of studies are neglected (Haddaway & Macura, 2018; Nuñez & Amano, 2021; Zenni et al., 2023). Multiple barriers to comprehensive evidence synthesis have been identified in previous global reviews and reports, such as the exclusion of grey literature (i.e., documents not published by commercial publishers) and non-English-language literature (Amano & Berdejo-Espinola, 2025; Chowdhury et al., 2022; Haddaway et al., 2015; Hannah et al., 2024; Nuñez & Amano, 2021; Pradier et al., 2026), the undervaluing of low impact factor journals (Choi et al., 2025), the transparency and availability of evidence sources (Dicks et al., 2014b; Hughes et al., 2024), and the underrepresentation of scientific evidence for certain species, communities and geographical regions (Amano et al., 2016; Christie et al., 2021; Hughes et al., 2023). It is already acknowledged that comprehensive evidence synthesis requires searching for non-English-language literature, grey literature and practitioner-held data (Berdejo-Espinola & Amano, 2025; Haddaway et al., 2015; Hannah et al., 2024; Nuñez & Amano, 2021), and that the extent of biases is arguably context-specific in influencing synthesis outcomes (Dobrescu et al., 2021; Nussbaumer-Streit et al., 2020). Nevertheless, despite ‘evidence-based conservation’ having been discussed for more than two decades, efforts to tackle systematic biases that can lead to poorly-informed decision-making in the conservation world remain limited, particularly in data-poor or underrepresented regions (Dicks et al., 2014b).

Assessing barriers to evidence-based conservation is particularly critical in Southeast Asia, a global centre of marine biodiversity facing severe and accelerating degradation from anthropogenic and climate pressures (Asaad et al., 2018; Kay et al., 2023), where the urgency for conservation action is high. Yet, the region’s information landscape poses unique challenges, with knowledge flowing across a mosaic of languages and ethnicities, scientific practices, and funding and publishing structures (Coleman et al., 2019). At least ten main languages are used in this region, with over 1,200 languages spoken across major and minor ethnic groups (Hammarström et al., 2025). Despite such linguistic diversity and the prevalence of grey literature, there are notably few literature reviews on ecology and conservation that incorporate multiple regional languages or extend beyond peer-reviewed sources. A single Southeast Asian study (Lim et al., 2024) has conducted a systematic search for peer-reviewed literature published in seven non-English regional languages to inform policymaking on marine plastic pollution. Frequently, reviews focusing on Southeast Asia narrow their English-language searches to specific countries rather than using regional languages to identify region-specific publications (e.g., Clark-Shen et al., 2023; Panyawai & Prathep 2022; and Stankovic et al., 2023). To date, no concerted effort has been made to include Southeast Asian languages and grey literature in evidence synthesis for this region, limiting the holistic integration of available knowledge into biodiversity assessment and applied conservation efforts.

Synthesising evidence for mobile and highly migratory species that rely on multiple habitats to complete critical life history stages presents particular challenges for conservation (Lascelles et al., 2014; Runge et al., 2014). Many undertake movements across the jurisdictions of multiple nations and management boundaries, which expose them to a compounding array of threats. Migratory species are also vulnerable to inadequate protection arising from the lack of coordinated international actions informed by comprehensive, spatially complete evidence of migratory connectivity – here defined as the degree to which areas within a landscape and seascape are linked by the movement or migration of individuals or populations (Harrison et al., 2018; Dunn et al., 2019; UNEP-WCMC, 2026). Knowledge gaps and biases in the underlying evidence base can misrepresent connectivity patterns, underestimate the importance of certain habitats, and misdirect limited conservation resources away from the populations and locations that require priority in protection (Boteler et al., 2022). Green turtles (*Chelonia mydas*), listed in the Appendices of the Convention on the Conservation of Migratory Species of Wild Animals (CMS), are an ideal model species for examining such biases. They demonstrate complex and extensive transboundary connectivity, and their movements are among the best documented for any marine taxon through the widespread use of animal-borne tracking technologies (Kot et al., 2022; Bentley et al., 2025; Restrepo et al., 2026). Southeast Asia supports some of the most important green turtle nesting and foraging aggregations globally, yet remains characterised as a data-poor region for connectivity research (Sequeira et al., 2025). This combination makes Southeast Asia a unique, relevant context in which to examine how gaps and biases in the available evidence base can affect the strength and reliability of information underpinning conservation decision making.

Here, we demonstrate substantial evidence on migratory connectivity for green turtles generated from Southeast Asia, which may not be included or utilised in meta-syntheses that systematically exclude grey literature and non-English-language literature. To examine this bias empirically, we conducted a comprehensive multi-language review of the movement of green turtles across Southeast Asia. Our objectives were to answer two principal questions: (1) *To what extent do limitations in publication types and languages bias the synthesis of migratory connectivity?* and (2) *How are green turtle migratory connections distributed and shared across national jurisdictions and regional management boundaries?* To address both objectives, we synthesised connectivity models at regional and national levels and quantified the effect of excluding grey literature and non-English-language sources on the resulting connectivity model. We also examined transboundary connectivity to inform international conservation policy, and assessed the delineation of existing Regional Management Units for green turtles across Southeast Asia. This information could help nations identify data gaps as well as counterpart countries whose cooperation is needed to improve transboundary conservation efforts. Ultimately, this review offers both a connectivity baseline to support monitoring and management, and a starting point for discussing strategies to overcome barriers and biases in synthesising new evidence, ensuring a more comprehensive understanding of the migratory connectivity of this species and the habitats they use.

## 2 Results

We gathered a comprehensive suite of literature across publications type and languages to develop a baseline of evidence on the movement and migratory connectivity of green turtles in Southeast Asia. To address selection and publication biases, we employed a systematic literature search method across global and regional bibliographic databases to retrieve both peer-reviewed and non-commercial grey literature, following the guidelines established by the Collaboration for Environmental Evidence (2013). Given that Southeast Asia is the focus of this review—where English is not the predominant language—the literature search was conducted in English and seven regional languages, specifically Filipino, Indonesian, Khmer, Malay, Thai, Traditional Chinese and Vietnamese.

### 2.1 Literature distribution

Using a combination of global and regional literature databases as well as unpublished data, we identified 115 references related to the movement and migration of green turtles. More than half (58%) of the references were grey literature, with an additional 8% being unpublished data, i.e. data not published in any form of report (**Figure 1**). Although unpublished data are sometimes distinguished from grey literature (Haddaway & Bayliss, 2015), we followed the broader definition of grey literature as material not formally published in peer-reviewed journals (Lefebvre et al., 2011; Haddaway et al., 2015) and treat both collectively as grey literature hereafter. Formally published literature which encompassed peer-reviewed journal articles and marine turtle-focused peer-reviewed newsletters only constituted a third of the references (34%). The vast majority of references (83%) were in English, while the remaining were in Traditional Chinese (11%), and in Indonesian, Vietnamese, Malay, and Thai (1-3%) (**Figure S1**). The geographic distribution of connectivity evidence was highly uneven, with Malaysia contributing the most references (n = 43) and Cambodia the fewest (n = 3) (**Table S3**). The disparity was further compounded by differences in publication effort, with institutions in Malaysia and Taiwan publishing the most (>20 references), while none of the publications came from Cambodian institutions (**Figure 2; Table S3**). Study and publishing institution locations did not always align. Specifically, a large portion of information from Thailand and Malaysia appeared in conference proceedings published in Japan, and most reports were published by regional intergovernmental organisations alongside national agencies, rather than universities or NGOs (**Table S3**).

**Figure 1.**
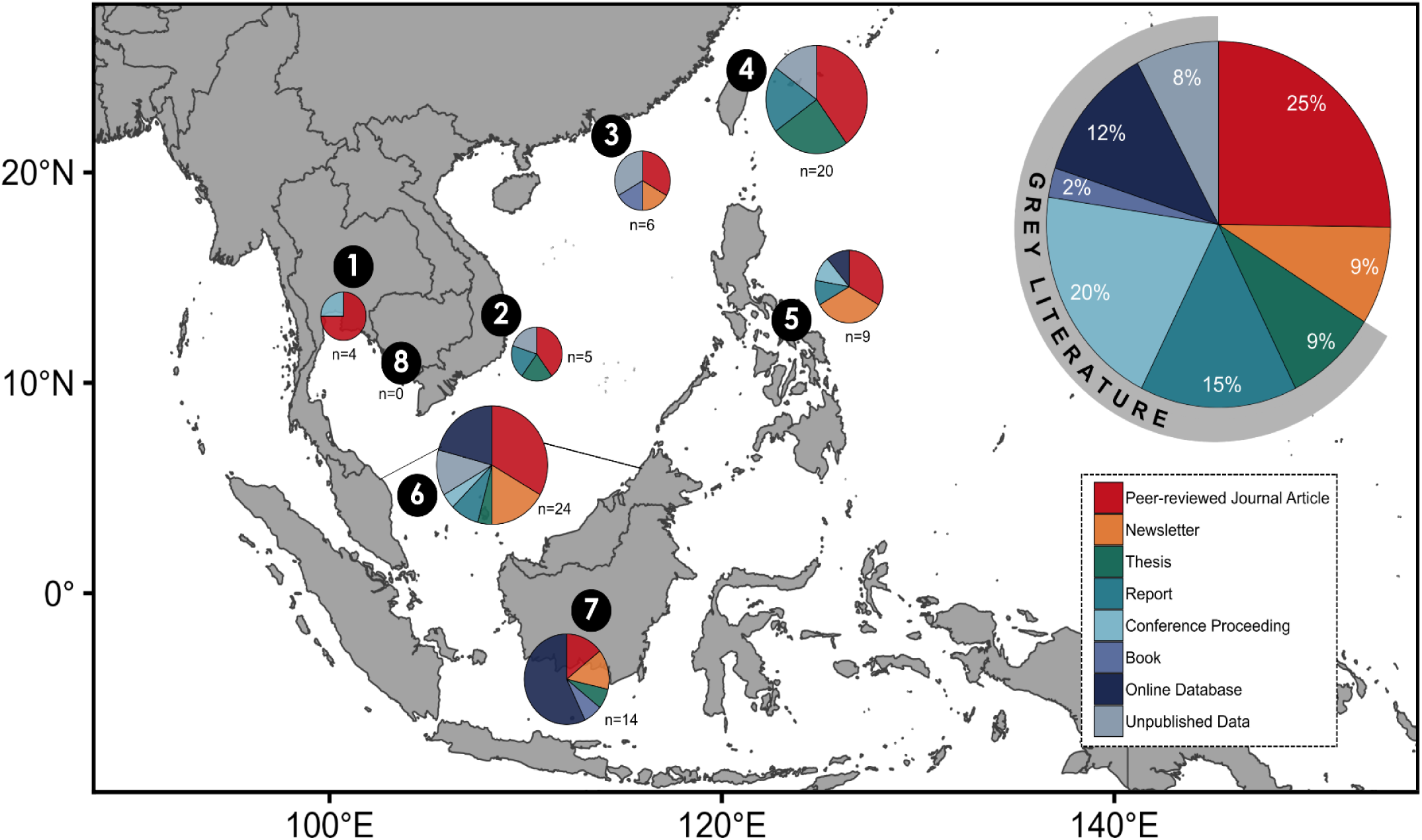
Literature types on green turtle movement published by each focal jurisdiction: (1) Thailand, (2) Vietnam, (3) Hong Kong, (4) Taiwan, (5) the Philippines, (6) Malaysia, and (7) Indonesia. No publications meeting the inclusion criteria were identified for (8) Cambodia. Pie size represents the total number of published literature included in this study. The larger pie chart shows the overall distribution of literature across literature types (n = 115).

**Figure 2.**
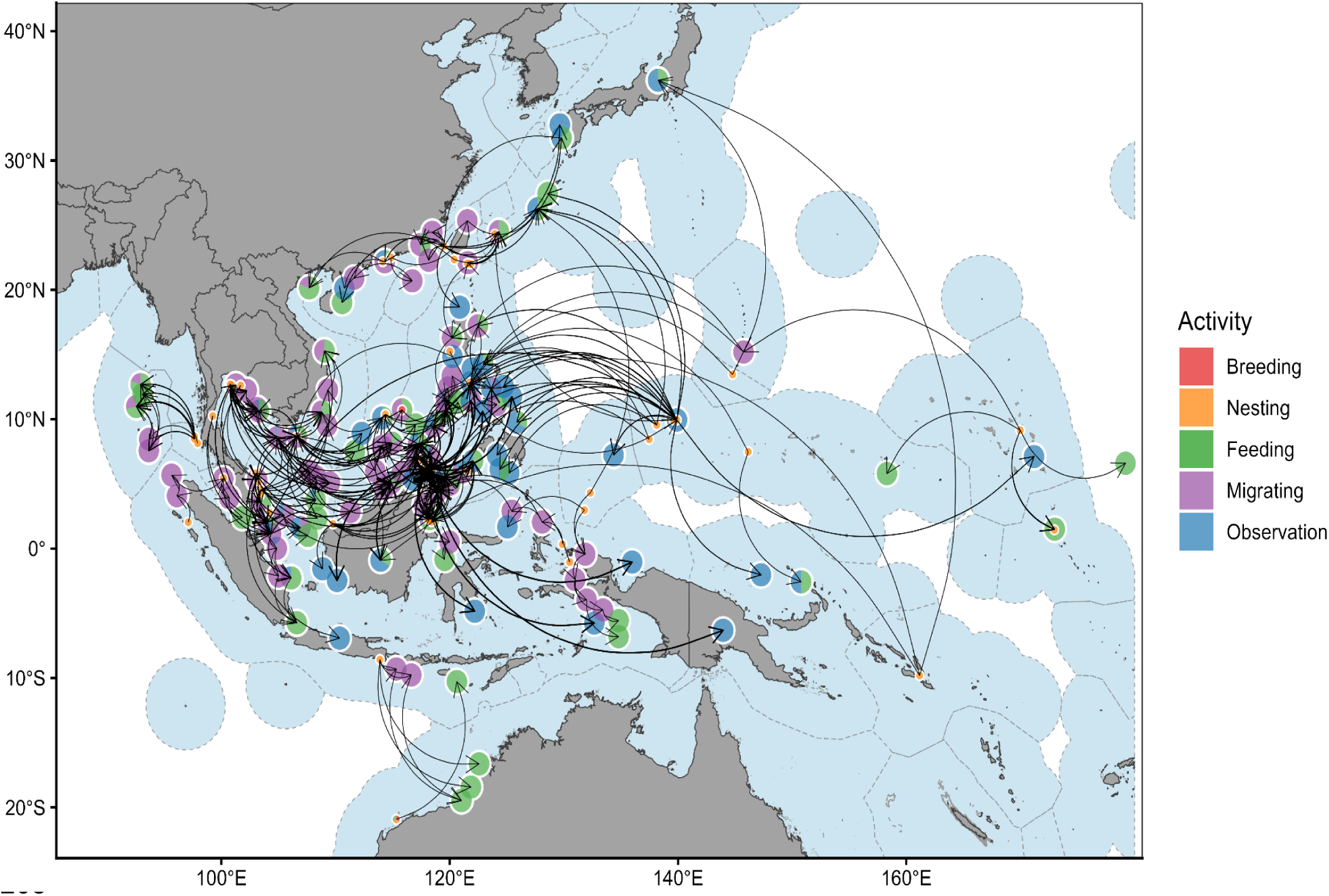
Transboundary connectivity of *Chelonia mydas* illustrating migrations, with nesting/breeding connected to feeding and non-feeding sites across EEZs of focal jurisdictions, including Cambodia, Hong Kong, Indonesia, Malaysia, the Philippines, Taiwan, Thailand, and Vietnam. Nodes, indicated by pie charts, represent metasites aggregated from individual sites, with segments showing the proportion of activity types at each metasite (i.e., breeding, nesting, feeding, migrating, and observation). Node size reflects site type, with a smaller radius (15 km) for reproductive sites (breeding and nesting) and a larger radius (100 km) for non-reproductive sites (feeding, migrating and observation). ‘Observation’ refers to those listed without a specifically documented behaviour or activity. Nodes on land are assigned to the EEZs of their respective jurisdictions without precise locations. Curved arrows indicate the movement directions, and line thickness reflects the scaled number of connections. Polygons around jurisdictions represent EEZs.

### 2.2 Transboundary movements across the EEZs and RMUs

We identified a total of 219 metasites (i.e., a site made up of one or more overlapping sites from different literature sources) and 416 metaconnections (ie., a connection between two sites made up of one or more connections found between those locations in different literature sources) for green turtles based on their individual movements (**Table 1**). Of these, 55 metasites were associated with nesting or breeding activities, while the remaining 164 were associated with non-reproductive activities, such as feeding or migrating. Given the diversity of political entities within the study area, including sovereign states, dependent territories, disputed territories, and special administrative regions, all are collectively referred to as ‘jurisdictions’ throughout this study for brevity. Every jurisdiction had at least one transboundary connection crossing into another jurisdiction’s Exclusive Economic Zone (EEZ) within Southeast Asia, East Asia or Oceania region (**Figure 2**). Metaconnection distances ranged from local-scale (5 km) to regional-scale (6,250 km), with an average of 862 km. We identified 242 large-scale metaconnections spanning >500 km across 22 jurisdictions, six of which are focal to this study: Indonesia, Malaysia, the Philippines, Taiwan, Thailand, and Vietnam.

**Table 1.**
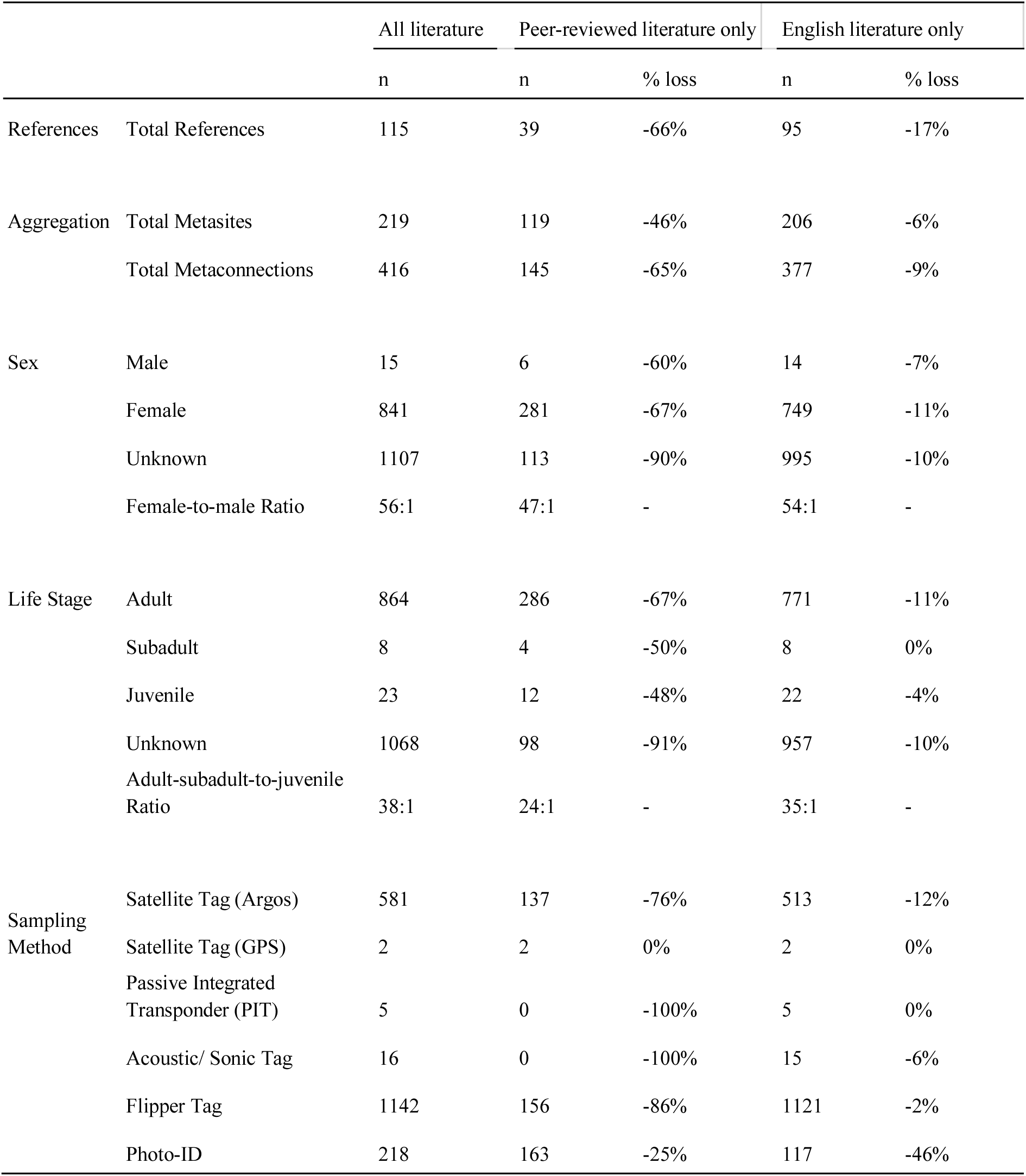
Summary of references assessing *Chelonia mydas* migratory connectivity across Southeast Asia, number of metasites and metaconnections, sampled individuals by sex and life stage, and sampling methods across three literature subsets: all literature, peer-reviewed literature only, and English literature only.

|  |  | All literature | Peer-reviewed literature only |  | English literature only |  |
| --- | --- | --- | --- | --- | --- | --- |
|  |  | n | n | % loss | n | % loss |
| References | Total References | 115 | 39 | -66% | 95 | -17% |
| Aggregation | Total Metasites | 219 | 119 | -46% | 206 | -6% |
|  | Total Metaconnections | 416 | 145 | -65% | 377 | -9% |
| Sex | Male | 15 | 6 | -60% | 14 | -7% |
|  | Female | 841 | 281 | -67% | 749 | -11% |
|  | Unknown | 1107 | 113 | -90% | 995 | -10% |
|  | Female-to-male Ratio | 56:1 | 47:1 | - | 54:1 | - |
| Life Stage | Adult | 864 | 286 | -67% | 771 | -11% |
|  | Subadult | 8 | 4 | -50% | 8 | 0% |
|  | Juvenile | 23 | 12 | -48% | 22 | -4% |
|  | Unknown | 1068 | 98 | -91% | 957 | -10% |
|  | Adult-subadult-to-juvenile Ratio | 38:1 | 24:1 | - | 35:1 | - |
| Sampling Method | Satellite Tag (Argos) | 581 | 137 | -76% | 513 | -12% |
|  | Satellite Tag (GPS) | 2 | 2 | 0% | 2 | 0% |
|  | Passive Integrated Transponder (PIT) | 5 | 0 | -100% | 5 | 0% |
|  | Acoustic/ Sonic Tag | 16 | 0 | -100% | 15 | -6% |
|  | Flipper Tag | 1142 | 156 | -86% | 1121 | -2% |
|  | Photo-ID | 218 | 163 | -25% | 117 | -46% |

Green turtles in the Philippines demonstrated the most integrated connectivity network, linking 14 jurisdictions (**Figure 3A; Table S4**). Most transboundary movement records (51 metaconnections) were recorded between Malaysia and the Philippines, particularly within and between sites in the Turtle Islands Heritage Protected Area (TIHPA), which comprises three Malaysian islands and six Philippine islands, as well as movements linking TIHPA to other sites within both jurisdictions. The green turtles in this area also exhibited extensive connectivity with Palau, Taiwan, Thailand, Papua New Guinea, and Indonesia. A large proportion of transboundary movements (30 metaconnections) were found between Malaysia (Peninsular, Sabah and Sarawak) and Indonesia, covering areas such as Bangka Island, Java, Southeast Sulawesi, West Kalimantan, and West Papua. No connectivity information related to turtles tracked from Cambodia or recaptured in Thailand was found in our literature review (**Figures S4 & S5**).

**Figure 3.**
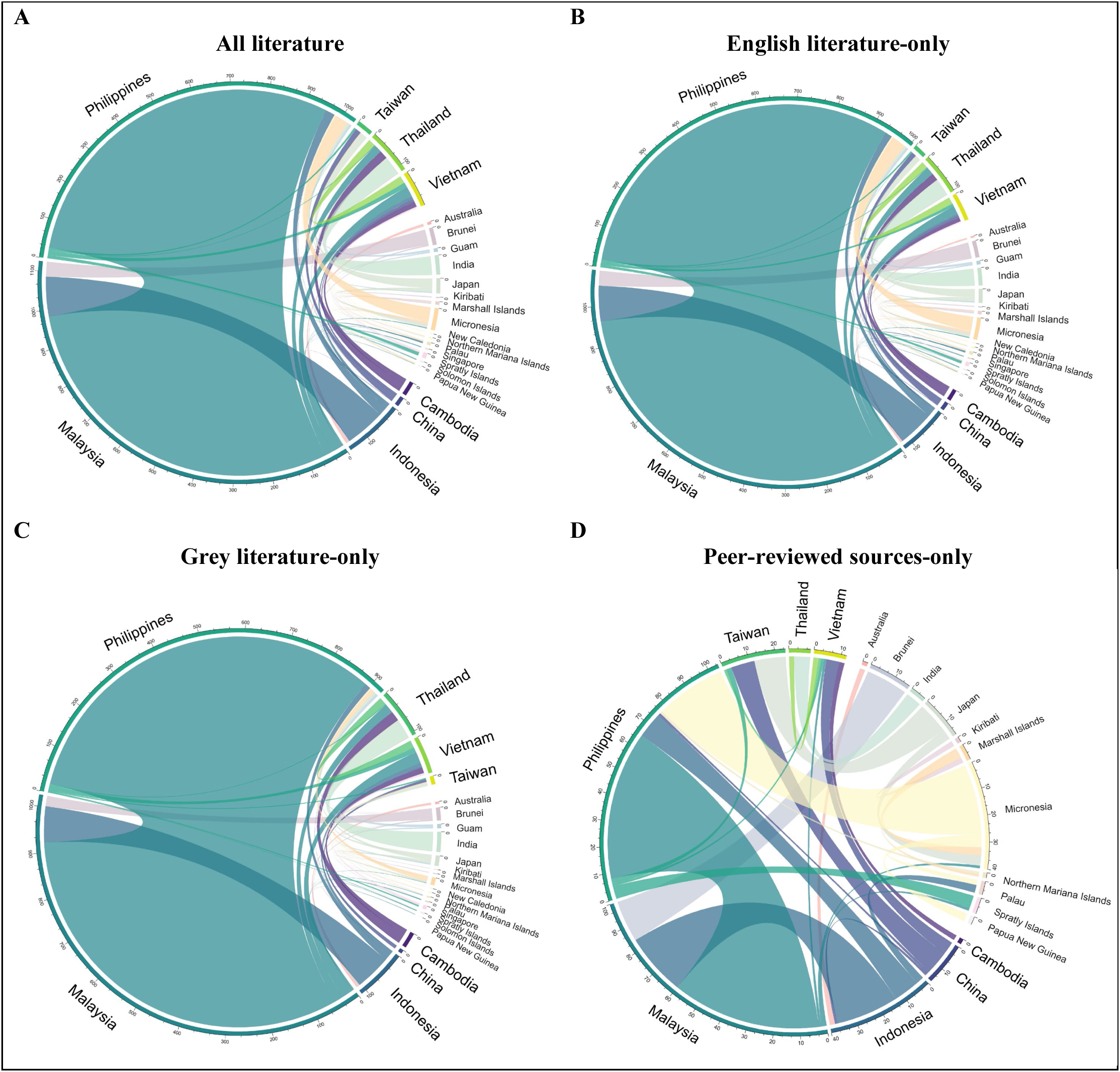
Connectivity of *Chelonia mydas* amongst jurisdictions, when (A) all literature, (B) only English literature, (C) only grey literature, and (D) only peer-reviewed sources, are included. The tick marks on the outer ring represent the number of individual-level connections to and from each jurisdiction, with the width of each chord proportional to the number of connections. Note that the directions of movement are not indicated.

Connectivity networks revealed that green turtles exhibit extensive movement within (91%) and across (9%) the boundaries of Regional Management Units (RMUs) (**Figure S7**). Notably, there were frequent crossings (32 metaconnections) between ‘East Indian and Southeast Asia (RMU 16)’ and ‘West Central Pacific (RMU 19)’. Additionally, connections were observed between the ‘Southwest Pacific (RMU 17)’ and both RMU 16 and RMU 19, including their overlapping areas, although these were less common (1-2 metaconnections).

### 2.3 Extent of language and publishing biases on an evidence base

#### 2.3.1 Impact of ignoring grey literature

Excluding grey literature resulted in substantial information loss across the green turtle connectivity network (**Table 1**; **Figure 4**). Nearly half of known metasites (46%; n = 100) and nearly two-thirds of known metaconnections (65%; n = 271) were lost. The impact was pervasive, where half of all metasites lost at least 88% of their documented connections when grey literature was excluded. At the jurisdiction level, the Philippines exhibited the highest connectivity in the full network, with 14 connected EEZs, and experienced the greatest loss upon grey literature exclusion, losing connections to 6 partner EEZs (representing 12% of the total country-level connections in the network; **Figure 3A & 3D, Table S4**). Malaysia, Thailand, and Indonesia each lost at least half of their country-level connections, while Taiwan (4 out of 5) and Vietnam (6 out of 8) retained most of their country-level connections in the peer-reviewed-only sources (**Table S4**).

**Figure 4.**
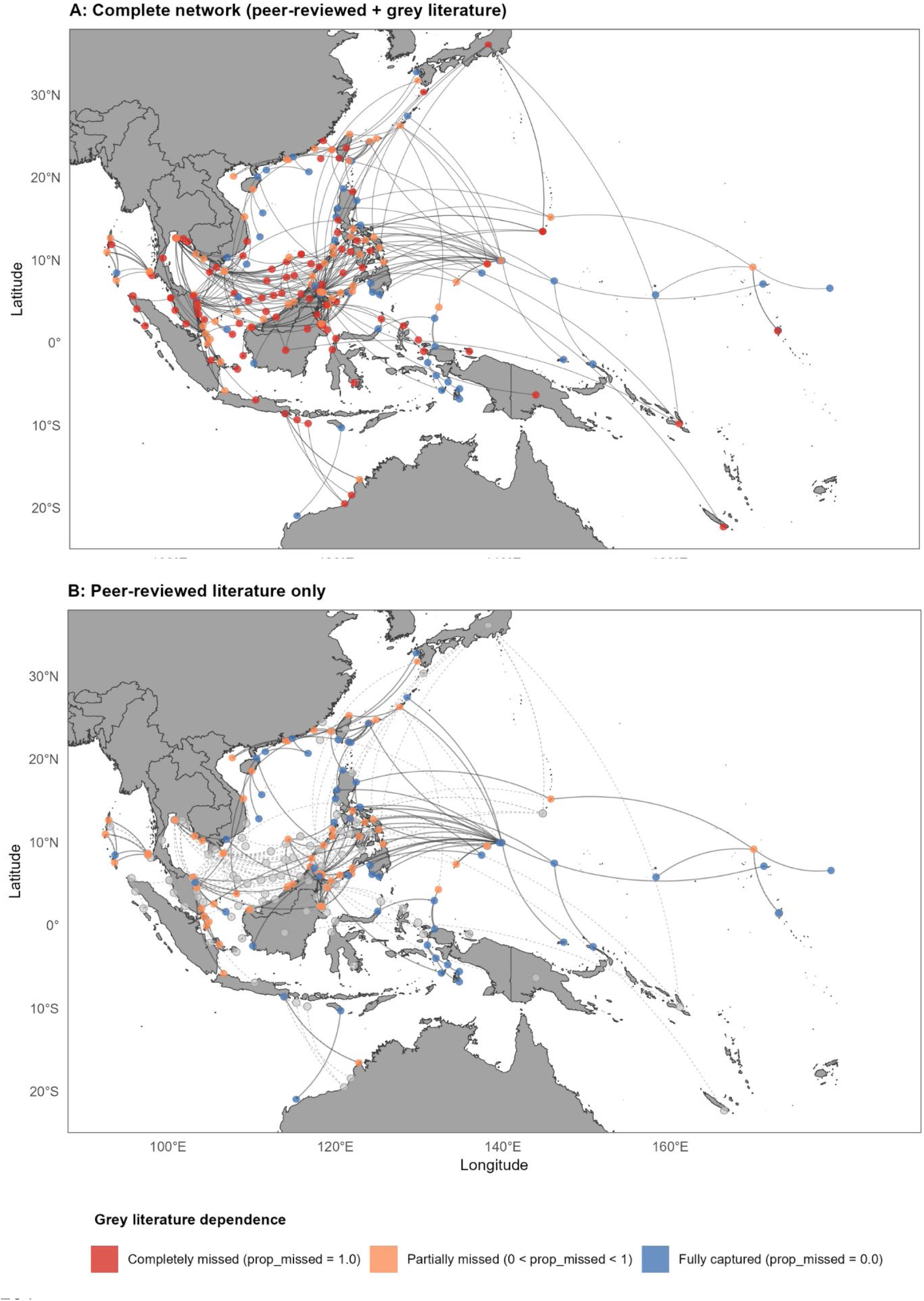
Regional connectivity network of *Chelonia mydas* illustrating (**A**) the complete network with all literature included, and (**B**) network using only peer-reviewed literature. Circles represent metasites and the lines represent metaconnections. The visible network is colour-coded based on grey literature dependency. Missed metasites and metaconnections resulting from grey literature exclusion are shown in grey.

Beyond connectivity, excluding grey literature also affected the demographic and methodological composition of the dataset. In particular, the apparent contribution of different sampling methods on the evidence-base shifted drastically when grey-literature was excluded. While the full dataset was dominated by flipper tags (n = 1,142) and satellite tags (n = 581), excluding grey literature removed 86% and 76% of the information from these two sampling methods, respectively, making photo-ID the dominant method in the subset of the dataset containing only peer-reviewed literature (n = 163, 75% retained). Demographic information of the green turtles was affected by the exclusion of grey literature; however, overall trends did not change. Particularly, male turtles decreased by 60%, and females by 67%, shifting the female-to-male ratio from 56:1 to 47:1, while the adult and subadult-to-juvenile-ratio decreased from 38:1 to 24:1. Finally, grey literature tended to have less metadata — its exclusion removed 90% of turtles with unknown sex and 91% of the turtles with unknown age classes, and a large proportion of observational records (those listed without specific behaviour or activity).

#### 2.3.2 Impact of ignoring non-English language literature

The effect of excluding non-English literature was much smaller than excluding grey literature. When non-English literature was excluded, the majority of metasites (68%) retained all documented metaconnections within English-language literature alone (**Table 1; Figure S3**). Non-English sources uniquely contributed to only 6% of all metasites (n = 13) that would otherwise be entirely undocumented. Both the spatial connectivity model (**Figure S3**) and the chord diagrams showing jurisdiction-to-jurisdiction connectivity (**Figure 3A & 3B**) showed very limited changes when generated from English language sources alone, indicating that metaconnections were mostly documented in English literature. All focal territories maintained their connected EEZs when considering only English literature, except Taiwan, which lost Micronesia among its connected countries (**Table S4**).

While the exclusion of grey literature artificially increased the apparent use of photo-ID relative to other sampling methods, excluding non-English sources overestimated the satellite telemetry usage compared to photo-ID, increasing from a ratio of 2.7:1 to 4.4:1 (**Table 1**). The effect of language restrictions was less notable on sex, life stage, and animal activity ratios.

## 3 Discussion

It is widely recognised that grey literature can contain substantial and vital evidence for applied conservation, and the reliability of a synthesis is undermined if it is based solely on a subset of available evidence (Bayliss and Beyer, 2014; CEE, 2013; Haddaway et al., 2015). Publication bias has been shown to affect at least 15-21% of meta-analyses of ecological studies (Jennions and Møller, 2002), yet many meta-analyses still fail to test for potential publication bias or do not have sufficient datasets for comprehensive analysis (Koricheva and Gurevitch, 2014). This study examined the impacts of systematic biases related to ignoring types of literature when developing an evidence base to inform conservation of a highly migratory species, the green turtle. We undertook an extensive multilingual search in eight languages, across peer-reviewed and grey literature. We developed the most comprehensive syntheses of green turtle movement and migratory connectivity in Southeast Asia. We quantified massive losses of information and shifts in the types of data that contribute to the evidence base resulting from the exclusion of grey literature, but less effect from the exclusion of non-English-language publications.

This review broadens the regional understanding of the migratory connectivity of green turtles, building on regional reviews compiled by Ng et al. (2018) and Pilcher et al. (2019), and a global review (Restrepo et al., 2026). From a transboundary conservation perspective, the regional- and jurisdiction-level connections described in this study (**Figure S4, S5, S6**) help elucidate responsibility for the management of migratory populations within and beyond national EEZs (Lascelles et al., 2014; Bentley et al., 2025). Our findings suggest that existing RMU delineations (Wallace et al., 2023) broadly capture the migratory patterns of green turtles in the region (91% of metaconnections occurring within RMU boundaries). The predominance of cross-boundary movements between the East Indian and Southeast Asia RMU (RMU 16) and the West Central Pacific RMU (RMU 19) is consistent with the latest IUCN subpopulation assessment (Pilcher, 2025), which treats these two RMUs as a single unit given the significant overlap and mixing of turtle populations between the two ocean basins.

Longstanding gaps in knowledge of species movements have been observed to limit the design and implementation of effective conservation strategies for highly mobile species, such as birds (Cottee-Jones et al., 2015; Scarpignato et al., 2023), elasmobranchs (Bortoluzzi et al., 2024; Bezerra et al., in review), and cetaceans (Murphy et al., 2021; Kot et al., 2023). Attempting to manage or conserve a migratory population such as sea turtles with incomplete information due to lack of consideration of grey literature can lead to erroneous or inefficient allocation of limited resources. For example, with the exclusion of grey literature, the Philippines would lose connectivity with six of its twelve partner countries–most of which are outside Southeast Asia, including Guam, Japan, New Caledonia, Palau, and the Solomon Islands. When looking across all connections, we found that at least half of all metasites would lose over 88% of their documented connections when grey literature is excluded. This represents a significant and avoidable loss of information that would undercut management and conservation.

Information on the migratory connectivity of green turtles in Southeast Asia is largely underpinned by grey literature, particularly ‘practitioner-generated research’ (Haddaway & Bayliss, 2015) such as conference proceedings and institutional reports. Conference proceedings constituted a large proportion of grey literature (30%), with a notable contribution from the Southeast Asia Sea Turtle Associative Research (SEASTAR2000) workshops (65% of all proceedings) between 2001 and 2011, accounting for the large proportion of Thai and Malaysian studies that appeared in proceedings published in Japan. Sea turtle tracking in this region is largely collaborative, involving government agencies, NGOs, universities, and local communities (Lau, 1999; Pilcher et al., 2019), reflected in the variety of report types that collectively constituted 22% of grey literature. Among these, government and intergovernmental reports dominated (82% of all reports), driven by national agency publishing efforts and the active role of the Southeast Asian Fisheries Development Center (SEAFDEC) in ASEAN sea turtle conservation since the 1990s.

While the importance of grey literature is obvious from their prevalence in the dataset and contributions to the connectivity models, their contribution is likely underestimated due to issues around information *accessibility*. While publishing reports and proceedings is a quicker and less costly way of getting information out compared to peer-reviewed journal articles, they can also be harder to access by policy-relevant bodies, particularly as they get older. For example, 41% of the reports included in this study could not be obtained through institutional websites, indicating that data-sharing policies or technical capacity could be a significant barrier to generating evidence bases to support management and conservation in the region. In particular, of the six government reports included in this study, only one was accessible on their respective agency website (**Figure S2**). Instead, they were obtained through direct communication with co-authors of this study or agency representatives. This difficulty in accessing governmental reports is underscored by the lack of any reports included in Thai, Indonesian and Khmer. To be clear, reports were known to exist in those languages, but due to inaccessibility it could not be determined whether they were relevant and should be included in our dataset.

The exclusion of non-English literature had a considerably smaller impact on the connectivity network compared to grey literature. Non-English sources made up 17% of the studies, but their exclusion from the connectivity model resulted in less than 10% of the metasites and metaconnections being removed. Previous ecological studies have shown that ignoring non-English literature can alter our understanding of the global environment, for instance by over-representating positive or statistically significant results, or by missing species- and habitat-specific information that exists only in the local language of non-English speaking countries (Amano et al., 2016; Konno et al., 2020; Amano et al., 2025). While omitting non-English references did not significantly change the connectivity network in this study, it influenced the characteristics of studies on regional green turtle tracking, where the reliance on satellite telemetry relative to photo-ID might be overestimated compared to actual practices. Although our findings suggest that excluding non-English publications did not alter the overall conclusions, understanding the underlying drivers of observed trends is crucial and warrants further investigation. The lack of references in Thai, Indonesian, and Khmer languages in our literature review may be due to accessibility issues, and that if these were accessible they could increase the impact of excluding non-English language sources (Hannah et al., 2025). Scientific publications and reports from the Philippines, and to some extent, Malaysia and Hong Kong, are mainly published in English, which explains the lack of non-English references from these regions.

While evidence of animal movement is consistently collected to varying degrees across jurisdictions, substantial information fails to be made accessible in a manner that allows for regional synthesis. The reasons include the reluctance of researchers, institutions, and governments to share datasets openly in public repositories and making published documents available online, and the lack of systematic and standardised management of biodiversity data (Scarpignato et al., 2023; Pastra et al., 2026). Monitoring data are not always published in academic journals due to several challenges, such as: (1) additional effort and resources are needed to analyse raw data to fit peer-review standards (Buxton et al., 2020; 2021); (2) insufficient data collected over meaningful timescales or with adequate sample sizes for robust conclusions (Lortie et al., 2007; Sunderland et al., 2009); or (3) telemetry studies that failed or were interrupted before completing the intended tracking period (Buxton et al., 2021). In this review, we found relatively few references of unpublished data, which were mainly contributed by conservation organisations and government agencies, as well as information sourced from social media or websites. However, it is likely that the number of references to unpublished data in our study is an underestimation of the true volume of unreported findings, likely due to how we structured the data collection of unpublished data (see Methods) as well as existing data sharing restrictions imposed on unpublished datasets.

From the findings of this review, our processes for gathering information to inform conservation need to be rethought. A multi-faceted (in terms of geography, language, publication type and institutions engaged) and multi-scale (from local to regional) approach is vital to develop unbiased evidence bases to support management and conservation. To bridge knowledge gaps effectively, collaborative efforts at a regional level are essential. This involves standardising and contributing data to online repositories (Dwyer et al., 2015; Rutz et al., 2020), reducing barriers to accessing governmental information (Hays et al., 2019; Driscoll et al., 2021; Södergren & Palm, 2021), and enhancing organisational capacity for the online indexing of documents containing data and knowledge (Tanalgo, 2025). These actions will enable us to synthesise a more comprehensive body of evidence, guiding the effective implementation of sea turtle conservation and management strategies.

Evidence-based approaches underpin global, cross-scale and transboundary policies (Segan et al., 2011; Sutherland et al., 2004). For migratory species in particular, addressing biases in evidence synthesis is crucial to support ongoing broad-scale and transboundary efforts like the ‘Important Area’ processes that synthesize data on different taxonomic groups with the intentions to make synthesised data accessible to marine conservation planners (e.g., Donald et al., 2019; Tetley et al., 2022; Kyne et al., 2023; Dunn et al., 2025; IUCN, 2016). Important Marine Turtle Areas (IMTAs) is the ‘Important Area’ process specifically for sea turtles. These ‘Important Area’ products are largely developed based on tracking data and expert opinion, both of which can be highly dependent on the information available in peer-reviewed literature. IMTAs, an emerging framework still under development (Bandimere et al., 2021), are a critical input into spatial planning, restoration prioritisation, and the development of networks of protected areas – thus they, and the evidence syntheses they are based on, directly inform the Kunming-Montreal Global Biodiversity Targets 1, 2 & 3 (respectively; CBD, 2022).

Implementing approaches that aim to develop a holistic and well-synthesised evidence base for tackling ecological gaps should be implemented across regions globally. Although this review primarily focuses on Southeast Asia, and adjacent regions including Taiwan and Hong Kong, the biases and barriers to knowledge extend beyond this region (Chowdhurry et al., 2022; Hughes et al., 2023; Hannah et al., 2025), regardless of development level or language spoken. Countries from the Global North are as likely to have community-based programs generating information that remains unpublished in peer-reviewed literature. In fact, all countries need mechanisms to appropriately aggregate and synthesise information from an array of sources to inform biodiversity reporting commitments (e.g., to the Convention on Biological Diversity or Convention on Migratory Species) in a comprehensive manner. If we do not address the methods used for evidence synthesis, we risk undermining the incredible resources and efforts currently focused on delivering these societal targets. What we protect will be shaped by what we missed in our efforts to synthesise vast sources of information.

## 4 Methods

### 4.1 Literature search

The literature search was conducted in English and seven regional languages, i.e. Filipino, Indonesian, Khmer, Malay, Thai, Traditional Chinese and Vietnamese, corresponding to the main languages used in the Philippines, Indonesia, Cambodia, Malaysia, Thailand, Hong Kong and Taiwan, and Vietnam. Ten reviewers searched the literature in their respective languages, with at least one regional database included for each non-English language (**Table S1**).

### 4.2 Search strategies

To systematically retrieve relevant literature, we developed English-language search strings based on previous search strategies used in marine megafauna studies (Kot et al., 2023a; Supplementary Material). The English search string was then translated into the seven regional languages. To evaluate the effectiveness, reviewers tested the translated search terms across different databases, particularly for terms with multiple possible translations. The most effective terms, i.e., with the greatest number of hits, were selected and incorporated into the final search strings. These were then adapted to fit the specific structures of each database. Whenever possible, searches were conducted within article titles, abstracts and keywords.

Initial scoping and backward searches, and direct communication with the lead author of a review on sea turtle tracking in Malaysia (Pilcher et al., 2019) revealed that a substantial amount of sea turtle movement data remains unpublished or is only accessible in reports unavailable through traditional literature searches. To address this limitation, additional searches were conducted through: (1) manually searching professional and governmental websites with relevance to the topic of systematic review (Haddaway & Bayliss, 2015; **Table S2**), (2) directly contacting researchers and organisations who were involved in collecting relevant data; (3) reviewing The State of the World’s Sea Turtles (SWOT) reports, and (4) reviewing publicly available social media information, e.g., Facebook, shared by contacted researchers and organisation members.

### 4.3 Data extraction and critical appraisal

Three reviewers were trained to extract data from full-text articles using standardised spreadsheets adapted from Kot et al. (2023a). The extracted information included: (i) general information, including publication year, publication type (peer-reviewed journal article, newsletter, thesis, report, conference proceeding, book, online database, and unpublished data), language, and study and publish location); (ii) site information (longitude and latitude of start and end locations); (iii) sampling method (satellite or acoustic telemetry, flipper tagging or photo-ID); (iv) individual information (number of individuals, species, sex, life stage, tracking date, and associated behaviour or activity); and (v) connectivity type (continuous track/route or point-to-point connection). For mark-recapture data from flipper tag and photo-ID studies, the study location was assigned to the jurisdiction where tags were deployed, as this reflects the sampling location and population origin. Recapture locations, which reflect realised spatial connectivity, were recorded separately.

To ensure the quality of the extracted data, we assessed the accuracy and the extent of the metadata provided in each study. Overall, we sought to be inclusive of varying levels of explicitness and uncertainty, taking into account the diversity in how information might be presented across peer-reviewed journals and grey literature. To ensure address the potential measurement bias based on unquantified uncertainty, each extracted piece of connectivity information was assigned a quality ranking based on the following simple but discriminatory criteria:

- Information was ranked as ‘low resolution’ and excluded from further analysis if any of the following features apply:

- *Unclear movement/sites*: Animal movement or connections between locations were difficult to interpret due to poor map resolution or a lack of explicit description.
- *Non-natural movement/sites*: Animal movement or connection between two locations resulted from translocated individuals rather than natural dispersal.
- Information was ranked as ‘medium resolution’ but was not excluded from the review if any of the following features apply:

- *Overly broad locations/sites*: Locations were described too generally (e.g., a country/territory name), potentially leading to biased site delineation.
- Information was ranked as ‘high resolution’ if none of the above limitations applied.

Each study was assessed by the reviewer responsible for data extraction. In cases of uncertainty, the final decision was made collectively by multiple reviewers. Reasons for exclusion were noted on the spreadsheet to ensure transparency.

### 4.4 Data analysis

The metadata of included references relating to the year of publication, jurisdiction (country/territory), language and type of document were summarised to understand the geographical and temporal distribution of the evidence on animal movement.

#### Creation of regional and sub-regional connectivity models

Connectivity models were synthesised at the regional and sub-regional (jurisdiction) levels. The models aimed to generate the most parsimonious network of sites and connections utilised by the species at scales that support policymakers, in addition to showing the breadth of work undertaken, published, and collated through this review process. The general approach to generating the connectivity models was adopted and modified from Dunn et al. (2019) and Bentley et al. (2025), who developed it to create a global dataset of migratory connectivity in the ocean (MiCO, www.mico.eco). For this study, individual georeferenced sites extracted from different studies were aggregated into ‘metasites’ through a three-step process: data preparation, site aggregation, and metasite creation. Details of each process are in the Supplementary Material. Final regional connectivity model outputs comprised points (metasites) and polylines (metaconnections) showing directions of migratory movements among Exclusive Economic Zones (EEZs; Flanders Marine Institute, 2024) and among marine turtle Regional Management Units (RMUs; Wallace et al., 2023), while sub-regional models showed migratory links between distinct sites within individual EEZs. The metaconnections that crossed EEZs were used to generate chord diagrams.

### Assessment of systematic biases from literature type and language exclusion

We assessed biases in how information on migratory connectivity of green turtles was published by quantifying the effect of excluding grey literature and non-English-language literature on the resulting connectivity model. For each metasite, we calculated the proportion of connections that would be lost if grey literature or non-English-language literature were excluded (prop_missed):

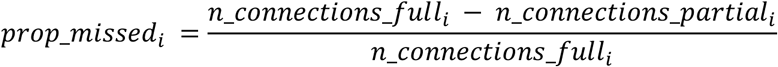

where *n_connections_fulli* is the number of unique connections documented at metasite *i* in the full dataset (peer-reviewed + grey literature; English + non-English), and *n_connections_partiali* is the number of connections documented in either peer-reviewed literature or English literature alone. A value of prop_missed = 0 indicates that all connections are fully captured by peer-reviewed or English literature, while prop_missed = 1 indicates that all connections are completely absent from peer-reviewed or English literature. This continuous metric reflects the gradient of systematic bias arising from reliance on grey literature and non-English-language sources. The spatial biases in the connectivity model derived from full and partial dataset were assessed within shared metasite boundaries to isolate the effect of literature and language type while controlling for spatial aggregation.

## Author contributions

Conceptualisation & design: HWKH, DCD

Literature search: HWKH, KCVB, TCC, NTD, REA, CLF, SMCH, DZH, SLL, CKYN, JP

Supervision: DCD

Formal analysis: HWKH, KCVB

Data extraction: HWKH, KCVB, DN

Writing—original draft: HWKH, DCD

Writing—review & editing: HWKH, KCVB, JR, TA, NP, CLF, DZH, SLL, CKYN, DCD

## References

Amano, T., & Berdejo-Espinola, V. (2025). Language barriers in conservation: consequences and solutions. Trends in Ecology & Evolution, 40(3), 273–285. 10.1016/j.tree.2024.11.003

Amano, T., Berdejo-Espinola, V., Christie, A. P., Willott, K., Akasaka, M., Baldi, A., Berthinussen, A., Bertolino, S., Bladon, A. J., Chen, M., Choi, C. Y., Kharrat, M. B. D., De Oliveira, L. G., Farhat, P., Golivets, M., Aranzamendi, N. H., Jantke, K., Kajzer-Bonk, J., Cisel Kemahli Aytekin, M., … Sutherland, W. J. (2021). Tapping into non-English-language science for the conservation of global biodiversity. PLoS Biology, 19(10). 10.1371/journal.pbio.3001296

Amano, T., Lamming, J. D. L., & Sutherland, W. J. (2016). Spatial Gaps in Global Biodiversity Information and the Role of Citizen Science. BioScience, 66(5), 393–400. 10.1093/biosci/biw022

Asaad, I., Lundquist, C. J., Erdmann, M. V., Van Hooidonk, R., & Costello, M. J. (2018). Designating Spatial Priorities for Marine Biodiversity Conservation in the Coral Triangle. Frontiers in Marine Science, 5. 10.3389/fmars.2018.00400

Bayliss, H. R., & Beyer, F. R. (2014). Information retrieval for ecological syntheses. Research Synthesis Methods, 6(2), 136–148. 10.1002/jrsm.1120

Bakker, E. S., Pagès, J. F., Arthur, R., & Alcoverro, T. (2016). Assessing the role of large herbivores in the structuring and functioning of freshwater and marine angiosperm ecosystems. Ecography, 39(2), 162–179. 10.1111/ecog.01651

Balvanera, P., Pascual, U., Christie, M., Baptiste, B., Lliso, B., Monroy, A.S., Guibrunet, L., Anderson, C.B., Athayde, S., Barton, D.N., Chaplin-Kramer, R., Jacobs, S., Kelemen, E., Kumar, R., Lazos, E., Martin, A., Mwampamba, T.H., Nakangu, B., O’Farrell, P., Raymond, C.M., Subramanian, S.M., Termansen, M., Van Noordwijk, M., Vatn, A., Contreras, V., and González-Jiménez, D. (2022). *Chapter 1: The role of the values of nature and valuation for addressing the biodiversity crisis and navigating towards more just and sustainable futures*. In: Methodological Assessment Report on the Diverse Values and Valuation of Nature of the Intergovernmental Science-Policy Platform on Biodiversity and Ecosystem Services. P. Balvanera, U. Pascual, C. Michael, B. Baptiste, and D. González-Jiménez (eds.). IPBES secretariat, Bonn, Germany. 10.5281/zenodo.6418971

Bandimere, A., Brenner, H., Casale, P., DiMatteo, A., Hurley, B., Hutchinson, B., Mast, R., Maxwell, S., Meyer, L., Poznik, Z., Rodrigues, I. & Wallace, B. (2021). Important Marine Turtle Areas Guidelines 1.0 (August 2021). Prepared for the 7th Burning Issues Workshop (BI-7), IUCN-SSC Marine Turtle Specialist Group. Available at: https://www.iucn-mtsg.org/s/IMTA-Guidelines-10.pdf [Accessed on 13 May 2026]

Bentley, L. K., Nisthar, D., Fujioka, E., Curtice, C., DeLand, S. E., Donnelly, B., Harrison, A.-L., Heywood, E. I., Kot, C. Y., Ortuño Crespo, G., Poulin, S., Halpin, P. N., & Dunn, D. C. (2025). Marine megavertebrate migrations connect the global ocean. Nature Communications, 16(1), 4089. 10.1038/s41467-025-59271-7

Berdejo-Espinola, V., & Amano, T. (2025). Assessing diverse values of nature requires multilingual evidence. Nature Reviews Biodiversity, 1(1), 5–6. 10.1038/s44358-024-00003-y

Bortoluzzi, J. R., McNicholas, G. E., Jackson, A. L., Klöcker, C. A., Ferter, K., Junge, C., Bjelland, O., Barnett, A., Gallagher, A. J., Hammerschlag, N., Roche, W. K., & Payne, N. L. (2024). Transboundary movements of porbeagle sharks support need for continued cooperative research and management approaches. Fisheries Research, 275, 107007. 10.1016/j.fishres.2024.107007

Boteler, B., Wagner, D., Durussel, C., Stokes, E., Gaymer, C. F., Friedlander, A. M., Dunn, D. C., Felipe Paredes Vargas, Véliz, D., & Hazin, C. (2022). Borderless conservation: Integrating connectivity into high seas conservation efforts for the Salas y Gómez and Nazca ridges. Frontiers in Marine Science, 9. 10.3389/fmars.2022.915983

Buxton, R. T., Avery-Gomm, S., Lin, H.-Y., Smith, P. A., Cooke, S. J., & Bennett, J. R. (2020). Half of resources in threatened species conservation plans are allocated to research and monitoring. Nature Communications, 11(1). 10.1038/s41467-020-18486-6

Buxton, R. T., Nyboer, E. A., Pigeon, K. E., Raby, G. D., Rytwinski, T., Gallagher, A. J., Schuster, R., Lin, H., Fahrig, L., Bennett, J. R., Cooke, S. J., & Roche, D. G. (2021). Avoiding wasted research resources in conservation science. Conservation Science and Practice, 3(2). 10.1111/csp2.329

CBD (Convention on Biological Diversity). (2022). Kunming-Montreal Global biodiversity framework. Available at: https://www.cbd.int/doc/c/e6d3/cd1d/daf663719a03902a9b116c34/cop-15-l-25-en.pdf [Accessed on 21 June 2026]

Ceballos, G., Ehrlich, P. R., & Dirzo, R. (2017). Biological annihilation via the ongoing sixth mass extinction signaled by vertebrate population losses and declines. Proceedings of the National Academy of Sciences of the United States of America, 114(30), E6089–E6096. 10.1073/pnas.1704949114

Choi, J. J., Gaskins, L. C., Morton, J. P., Bingham, J. A., Blawas, A. M., Hayes, C., Hoyt, C., Halpin, P. N., & Silliman, B. (2025). Role of low-impact-factor journals in conservation implementation. Conservation Biology, 39(2), e14391. 10.1111/cobi.14391

Chowdhury, S., Gonzalez, K., Aytekin, M. Ç. K., Baek, S. Y., Bełcik, M., Bertolino, S., Duijns, S., Han, Y., Jantke, K., Katayose, R., Lin, M. M., Nourani, E., Ramos, D. L., Rouyer, M. M., Sidemo-Holm, W., Vozykova, S., Zamora-Gutierrez, V., & Amano, T. (2022). Growth of non-English-language literature on biodiversity conservation. Conservation Biology, 36(4). 10.1111/cobi.13883

Christie, A. P., Amano, T., Martin, P. A., Petrovan, S. O., Shackelford, G. E., Simmons, B. I., Smith, R. K., Williams, D. R., Wordley, C. F. R., & Sutherland, W. J. (2021). The challenge of biased evidence in conservation. Conservation Biology, 35(1), 249–262. 10.1111/cobi.13577

Clark-Shen, N., Chin, A., Arunrugstichai, S., Labaja, J., Mizrahi, M., Simeon, B., & Hutchinson, N. (2023). Status of Southeast Asia’s marine sharks and rays. Conservation Biology, 37(1). 10.1111/cobi.13962

Coleman, J. L., Ascher, J. S., Bickford, D., Buchori, D., Cabanban, A., Chisholm, R. A., Chong, K. Y., Christie, P., Clements, G. R., dela Cruz, T. E. E., Dressler, W., Edwards, D. P., Francis, C. M., Friess, D. A., Giam, X., Gibson, L., Huang, D., Hughes, A. C., Jaafar, Z., Jain, A… & Carrasco, L. R. (2019). Top 100 research questions for biodiversity conservation in Southeast Asia. Biological Conservation, 234, 211–220. 10.1016/j.biocon.2019.03.028

Collaboration for Environmental Evidence, CEE. (2013). Guidelines for Systematic Review and Evidence Synthesis in Environmental Management. Version 4.2. www.environmentalevidence.org/Documents/Guidelines/Guidelines4.2.pdf

Cook, C. N., Carter, R. W. B., Fuller, R. A., & Hockings, M. (2012). Managers consider multiple lines of evidence important for biodiversity management decisions. Journal of Environmental Management, 113, 341–346. 10.1016/j.jenvman.2012.09.002

Dicks, L. V., Hodge, I., Randall, N. P., Scharlemann, J. P. W., Siriwardena, G. M., Smith, H. G., Smith, R. K., & Sutherland, W. J. (2014a). A Transparent Process for “Evidence-Informed” Policy Making. Conservation Letters, 7(2), 119–125. 10.1111/conl.12046

Dicks, L. V., Walsh, J. C., & Sutherland, W. J. (2014b). Organising evidence for environmental management decisions: A “4S” hierarchy. In Trends in Ecology and Evolution (Vol. 29, Number 11, pp. 607–613). Elsevier Ltd. 10.1016/j.tree.2014.09.004

Dobrescu, A., Nussbaumer-Streit, B., Klerings, I., Wagner, G., Persad, E., Sommer, I., Herkner, H., & Gartlehner, G. (2021). Restricting evidence syntheses of interventions to English-language publications is a viable methodological shortcut for most medical topics: a systematic review. Journal of Clinical Epidemiology, 137, 209–217. 10.1016/j.jclinepi.2021.04.012

Donald, P.F., Fishpool, L.D., Ajagbe, A., Bennun, L.A., Bunting, G., Burfield, I.J., Butchart, S.H., Capellan, S., Crosby, M.J., Dias, M.P. and Diaz, D., 2019. Important Bird and Biodiversity Areas (IBAs): the development and characteristics of a global inventory of key sites for biodiversity. Bird Conservation International, 29(2), pp.177–198.

Driscoll, D. A., Garrard, G. E., Kusmanoff, A. M., Dovers, S., Maron, M., Preece, N., Pressey, R. L., & Ritchie, E. G. (2020). Consequences of information suppression in ecological and conservation sciences. Conservation Letters, 14(1). 10.1111/conl.12757

Dunn, D. C., Harrison, A. L., Curtice, C., DeLand, S., Donnelly, B., Fujioka, E., Heywood, E., Kot, C. Y., Poulin, S., Whitten, M., Åkesson, S., Alberini, A., Appeltans, W., Arcos, J. M., Bailey, H., Ballance, L. T., Block, B., Blondin, H., Boustany, A. M., … Halpin, P. N. (2019). The importance of migratory connectivity for global ocean policy. Proceedings of the Royal Society B: Biological Sciences, 286(1911). 10.1098/rspb.2019.1472

Dunn, D. C., Cleary, J., DeLand, S., Bax, N., Bentley, L. K., Curtice, C., Donnelly, B., Dunstan, P. K., C. Barrio Froján, Gjerde, K. M., Gunn, V., Johnson, D. E., Klein, E., Kot, C. Y., D. Nisthar, Crespo, G. O., & Halpin, P. N. (2025). What is an ecologically or biologically significant area? Npj Ocean Sustainability, 4(1). 10.1038/s44183-025-00126-5

Fraisl, D., Campbell, J., See, L., Wehn, U., Wardlaw, J., Gold, M., Moorthy, I., Arias, R., Piera, J., Oliver, J. L., Masó, J., Penker, M., & Fritz, S. (2020). Mapping citizen science contributions to the UN sustainable development goals. Sustainability Science 2020 15:6, 15(6), 1735–1751. 10.1007/s11625-020-00833-7

Godley, B., Blumenthal, J., Broderick, A., Coyne, M., Godfrey, M., Hawkes, L., & Witt, M. (2008). Satellite tracking of sea turtles: Where have we been and where do we go next? Endangered Species Research, 4, 3–22. 10.3354/esr00060

Haddaway, N. R., & Bayliss, H. R. (2015). Shades of grey: Two forms of grey literature important for reviews in conservation. Biological Conservation, 191, 827–829. 10.1016/j.biocon.2015.08.018

Haddaway, N. R., Collins, A. M., Coughlin, D., & Kirk, S. (2015). The role of google scholar in evidence reviews and its applicability to grey literature searching. PLoS ONE, 10(9). 10.1371/journal.pone.0138237

Haddaway, N. R., & Macura, B. (2018). The role of reporting standards in producing robust literature reviews. Nature Climate Change 2018 8:6, 8(6), 444–447. 10.1038/s41558-018-0180-3

Halpin, P.N., A.J. Read, E. Fujioka, B.D. Best, B. Donnelly, L.J. Hazen, C. Kot, K. Urian, E. LaBrecque, A. Dimatteo, J. Cleary, C. Good, L.B. Crowder, and K.D. Hyrenbach. (2009). OBIS-SEAMAP: The world data center for marine mammal, sea bird, and sea turtle distributions. Oceanography 22(2):104–115

Hammarström, H., et al. (2025). Glottolog 5.0. Leipzig: Max Planck Institute for Evolutionary Anthropology.

Hannah, K., Fuller, R. A., Smith, R. K., Sutherland, W. J., & Amano, T. (2025). Language barriers in conservation science citation networks. Conservation Biology. 10.1111/cobi.70051

Hannah, K., Haddaway, N. R., Fuller, R. A., & Amano, T. (2024). Language inclusion in ecological systematic reviews and maps: Barriers and perspectives. Research Synthesis Methods, 15(3), 466–482. 10.1002/jrsm.1699

Harrison, A. L., Costa, D. P., Winship, A. J., Benson, S. R., Bograd, S. J., Antolos, M., Carlisle, A. B., Dewar, H., Dutton, P. H., Jorgensen, S. J., Kohin, S., Mate, B. R., Robinson, P. W., Schaefer, K. M., Shaffer, S. A., Shillinger, G. L., Simmons, S. E., Weng, K. C., Gjerde, K. M., & Block, B. A. (2018). The political biogeography of migratory marine predators. Nature Ecology and Evolution, 2(10), 1571–1578. 10.1038/s41559-018-0646-8

Hays, G. C., & Hawkes, L. A. (2018). Satellite Tracking Sea Turtles: Opportunities and Challenges to Address Key Questions. Frontiers in Marine Science, 5. 10.3389/fmars.2018.00432

Hays, G. C., Bailey, H., Bograd, S. J., Bowen, W. D., Campagna, C., Carmichael, R. H., Casale, P., Chiaradia, A., Costa, D. P., Cuevas, E., Nico de Bruyn, P. J., Dias, M. P., Duarte, C. M., Dunn, D. C., Dutton, P. H., Esteban, N., Friedlaender, A., Goetz, K. T., Godley, B. J., & Halpin, P. N. (2019). Translating Marine Animal Tracking Data into Conservation Policy and Management. Trends in Ecology & Evolution, 34(5), 459–473. 10.1016/j.tree.2019.01.009

Hill, J. T., Olds, A. D., Gilby, B. L., Mosman, J. D., James, A. L., & Henderson, C. J. (2025). Seascape connectivity shapes fish functional diversity, ecosystem functioning and resilience in subtropical coral reefs. Coral Reefs. 10.1007/s00338-025-02734-6

Hudson, L. N., Newbold, T., Contu, S., Hill, S. L. L., Lysenko, I., De Palma, A., Phillips, H. R. P., Alhusseini, T. I., Bedford, F. E., Bennett, D. J., Booth, H., Burton, V. J., Chng, C. W. T., Choimes, A., Correia, D. L. P., Day, J., Echeverría-Londoño, S., Emerson, S. R., Gao, D., … Purvis, A. (2017). The database of the PREDICTS (Projecting Responses of Ecological Diversity In Changing Terrestrial Systems) project. Ecology and Evolution, 7(1), 145–188. 10.1002/ece3.2579

Hughes, A. C., Than, K. Z., Tanalgo, K. C., Agung, A. P., Alexander, T., Kane, Y., Bhadra, S., Chornelia, A., Sritongchuay, T., Simla, P., Chen, Y., Chen, X., Uddin, N., Khatri, P., & Karlsson, C. (2023). Who is publishing in ecology and evolution? the underrepresentation of women and the Global South. Frontiers in Environmental Science, 11. 10.3389/fenvs.2023.1211211

Hughes, A., Orr, M., Palacio, R., Xuan, Y., & Qiao, H. (2024). A dire need for better standards of data quality, transparency, and reproducibility in IUCN RedList assessments. 10.32942/X2KK7D

IPBES. (2019). Global assessment report on biodiversity and ecosystem services of the Intergovernmental Science-Policy Platform on Biodiversity and Ecosystem Services. 10.5281/zenodo.6417333

IUCN. (2016). A Global Standard for the Identification of Key Biodiversity Areas, Version 1.0. First edition. Gland, Switzerland: IUCN.

Jennions, M. D., & Møller, A. P. (2002). Publication bias in ecology and evolution: an empirical assessment using the “trim and fill” method. Biological Reviews of the Cambridge Philosophical Society, 77(2), 211–222. 10.1017/s1464793101005875

Kay, S., Avillanosa, A. L., Cheung, V., Dao, H. N., Gonzales, B. J., Palla, H. P., Praptiwi, R. A., Queirós, A. M., Sailley, S. F., Sumeldan, J., Wan Mohd Syazwan, Then, A. Y-H, & Wee, H. B. (2023). Projected effects of climate change on marine ecosystems in Southeast Asian seas. Frontiers in Marine Science, 10. 10.3389/fmars.2023.1082170

Koricheva, J., & Gurevitch, J. (2014). Uses and misuses of meta-analysis in plant ecology. Journal of Ecology, 102(4), 828–844. 10.1111/1365-2745.12224

Kot, C. Y., DeLand, S. E., Harrison, A.-L., Alberini, A., Blondin, H., Chory, M., Cleary, J., Curtice, C., Donnelly, B., Fujioka, E., Palacio, A. H., Heywood, E. I., Mason, E., Nisthar, D., Crespo, G. O., Poulin, S., Whitten, M., Woolston, C., Dunn, D. C., & Halpin, P. N. (2023a). Synthesizing connectivity information from migratory marine species for area-based management. Biological Conservation, 283, 110142. 10.1016/j.biocon.2023.110142

Kot, C.Y., E. Fujioka, A.D. DiMatteo, A. Bandimere, B.P. Wallace, B.J. Hutchinson, J. Cleary, P.N. Halpin and R.B. Mast. (2023b). The State of the World’s Sea Turtles Online Database: Data provided by the SWOT Team and hosted on OBIS-SEAMAP. Oceanic Society, Conservation International, IUCN Marine Turtle Specialist Group (MTSG), and Marine Geospatial Ecology Lab, Duke University. http://seamap.env.duke.edu/swot

Kot, C. Y., Åkesson, S., Alfaro-Shigueto, J., Llanos, D. F. A., Antonopoulou, M., Balazs, G. H., Baverstock, W. R., Blumenthal, J. M., Broderick, A. C., Bruno, I., Canbolat, A. F., Casale, P., Cejudo, D., Coyne, M. S., Curtice, C., DeLand, S., DiMatteo, A., Dodge, K., Dunn, D. C., … Halpin, P. N. (2022). Network analysis of sea turtle movements and connectivity: A tool for conservation prioritization. Diversity and Distributions, 28(4), 810–829. 10.1111/ddi.13485

Kyne, P.M., di Sciara, G.N., Morera, A.B., Charles, R., Rodríguez, E.G., Fernando, D., Pestana, A.G., Priest, M. and Jabado, R.W., 2023. Important Shark and Ray Areas: a new tool to optimize spatial planning for sharks. Oryx, 57(2), pp.146–147.

Lascelles, B., Notarbartolo Di Sciara, G., Agardy, T., Cuttelod, A., Eckert, S., Glowka, L., Hoyt, E., Llewellyn, F., Louzao, M., Ridoux, V., & Tetley, M. J. (2014). Migratory marine species: Their status, threats and conservation management needs. Aquatic Conservation: Marine and Freshwater Ecosystems, 24(S2), 111–127. 10.1002/aqc.2512

Lefebvre, C., Manheimer, E., Glanville, J. Chapter 6: Searching for studies. In: Higgins, J. P., Green, S., editor(s). Cochrane Handbook for Systematic Reviews of Interventions Version 5.1.0 (updated March 2011). The Cochrane Collaboration, 2011. Available from: https://training.cochrane.org/versions-and-changes-handbook

Lim, C. L., Lyons, Y., Liu, Y., Neo, M. L., Müller, M., Wong, C., Cordova, M. R., Sulistiowati, S., Abreo, N. A. S., Ko Gyi, T., Onda, D. F., Baculi, R., Charoenpong, C., Lalung, J., Le Hoang, H. A., Li, D., & Zhu, L. (2024). Literature-based database to inform policy making on marine plastic pollution in ASEAN+3. Frontiers in Ocean Sustainability, 2. 10.3389/focsu.2024.1356148

Lortie, C. J., Aarssen, L. W., Budden, A. E., Koricheva, J., Roosa Leimu, & Tregenza, T. (2007). Publication bias and merit in ecology. Oikos, 116(7), 1247–1253. 10.1111/j.2007.0030-1299.15686.x

Flanders Marine Institute. (2024). Marineregions: intersect of EEZs and IHO areas (Version 5 - 2024). Available online at https://www.marineregions.org/.

McLean, M., Warner, B., Markham, R., Fischer, M., Walker, J., Klein, C., Hoeberechts, M., & Dunn, D. C. (2023). Connecting conservation & culture: The importance of Indigenous Knowledge in conservation decision-making and resource management of migratory marine species. Marine Policy, 155(3), 105582. 10.1016/j.marpol.2023.105582

Mumby, P. J., Edwards, A. J., Arias-González, J. E., Lindeman, K. C., Blackwell, P. G., Gall, A., Gorczynska, M. I., Harborne, A. R., Pescod, C. L., Renken, H., Wabnitz, C. C. C., & Llewenyn, G. (2004). Mangroves enhance the biomass of coral reef fish communities in the Caribbean. Nature, 427(6974), 533–536. 10.1038/nature02286

Murphy, E. J., Johnston, N. M., Hofmann, E. E., Phillips, R. A., Jackson, J. A., Constable, A. J., Henley, S. F., Melbourne-Thomas, J., Trebilco, R., Cavanagh, R. D., Tarling, G. A., Saunders, R. A., Barnes, D. K. A., Costa, D. P., Corney, S. P., Fraser, C. I., Höfer, J., Hughes, K. A., Sands, C. J., & Thorpe, S. E. (2021). Global Connectivity of Southern Ocean Ecosystems. Frontiers in Ecology and Evolution, 9. 10.3389/fevo.2021.624451

Nuñez, M. A., & Amano, T. (2021). Monolingual searches can limit and bias results in global literature reviews. Nature Ecology & Evolution, 5(3), 264–264. 10.1038/s41559-020-01369-w

Nuñez, M. A., Chiuffo, M. C., Pauchard, A., & Zenni, R. D. (2021). Making ecology really global. Trends in Ecology & Evolution. 10.1016/j.tree.2021.06.004

Nussbaumer-Streit, B., Klerings, I., Dobrescu, A. I., Persad, E., Stevens, A., Garritty, C., Kamel, C., Affengruber, L., King, V. J., & Gartlehner, G. (2020). Excluding non-English publications from evidence-syntheses did not change conclusions: a meta-epidemiological study. Journal of Clinical Epidemiology, 118, 42–54. 10.1016/j.jclinepi.2019.10.011

Panyawai, J., & Prathep, A. (2022). A Systematic Review of the Status, Knowledge, and Research Gaps of Dugong in Southeast Asia. Aquatic Mammals, 48, 203–222. 10.1578/AM.48.3.2022.203

Pastra, A., Muller-Karger, F. E. & Soares, J. (2026). Data Challenges in the Observation of Marine Biodiversity: Toward a Collaborative Framework. In Kraska, J., Long, R. & Pedrozo, R. (Eds.), Challenges in oceans law and policy: The South Pacific and Latin America (Vol. 26, pp. 50–75). *Brill*. 10.1163/9789004760806

Perez, M. A., Limpus, C. J., Hofmeister, K., Shimada, T., Strydom, A., Webster, E., & Hamann, M. (2022). Satellite tagging and flipper tag recoveries reveal migration patterns and foraging distribution of loggerhead sea turtles (*Caretta caretta*) from eastern Australia. Marine Biology, 169(6). 10.1007/s00227-022-04061-8

Pilcher, N.J. 2025. *Chelonia mydas* (East Indian–West Pacific subpopulation). The IUCN Red List of Threatened Species 2025: e.T220970168A220970208. 10.2305/IUCN.UK.2025-2.RLTS.T220970168A220970208.en. [Accessed on 22 June 2026]

Pilcher, N. J., Bali, J., Buis, J., Eng Heng, C., Devadasan, A., Isnain, I., Haniza Binti Jamil, N., Joseph, J., Min Min, L., Hock Chark, L., Abdullah Bin Syed Abdul Kadir, S., Ruqaiyah, S., Bracken Tisen, O., Van De Merwe, J. P., & Williams, J. (2019). A REVIEW OF SEA TURTLE SATELLITE TRACKING IN MALAYSIA. Indian Ocean Turtle Newsletter, (29), 11–22.

Pradier, C., Céspedes, L., & Larivière, V. (2026). How multilingual is scholarly communication? Mapping the global distribution of languages in publications and citations. Journal of the Association for Information Science and Technology. 10.1002/asi.70055

Pullin, A. S., & Knight, T. M. (2012). Science informing Policy – a health warning for the environment. Environmental Evidence 2012 1:1, 1(1), 15-. 10.1186/2047-2382-1-15

Restrepo, J., Nisthar, D., Heng, H. W., Bentley, L. K., Curtice, C., DeLand, S., Fujioka, E., Halpin, P. N., Poulin, S. K., Richardson, A. J., Seminoff, J. A., Valverde, R. A., & Dunn, D. C. (2026). Ecological Insights and Management Implications of the Global Migratory Connectivity of Green Turtles. Diversity and Distributions, 32(5). 10.1111/ddi.70196

Reyes-García, V., García-del-Amo, D., Álvarez-Fernández, S., Benyei, P., Calvet-Mir, L., Junqueira, A. B., Labeyrie, V., Li, X., Miñarro, S., Porcher, V., Porcuna-Ferrer, A., Schlingmann, A., Schunko, C., Soleymani, R., Tofighi-Niaki, A., Abazeri, M., Attoh, E. M. N. A. N., Ayanlade, A., Ávila, J. V. D. C., … Zakari, I. S. (2024). Indigenous Peoples and local communities report ongoing and widespread climate change impacts on local social-ecological systems. Communications Earth & Environment 2024 5:1, 5(1), 29-. 10.1038/s43247-023-01164-y

Runge, C. A., Martin, T. G., Possingham, H. P., Willis, S. G., & Fuller, R. A. (2014). Conserving mobile species. Frontiers in Ecology and the Environment, 12(7), 395–402. 10.1890/130237

Rutz, C., Loretto, M.-C., Bates, A. E., Davidson, S. C., Duarte, C. M., Jetz, W., Johnson, M., Kato, A., Kays, R., Mueller, T., Primack, R. B., Ropert-Coudert, Y., Tucker, M. A., Wikelski, M., & Cagnacci, F. (2020). COVID-19 lockdown allows researchers to quantify the effects of human activity on wildlife. Nature Ecology & Evolution, 4. 10.1038/s41559-020-1237-z

Scarpignato, A. L., Huysman, A. E., Jimenez, M. F., Witko, C. J., Harrison, A.-L., Seavy, N. E., Smith, M. A., Deppe, J. L., Wilsey, C. B., & Marra, P. P. (2023). Shortfalls in tracking data available to inform North American migratory bird conservation. Biological Conservation, 286, 110224. 10.1016/j.biocon.2023.110224

Segan, D. B., Bottrill, M. C., Baxter, P. W. J., & Possingham, H. P. (2011). Using conservation evidence to guide management. Conservation Biology, 25(1), 200–202. 10.1111/j.1523-1739.2010.01582.x

Sequeira, A. M., Rodríguez, J. P., Marley, S. A., Calich, H. J., van der Mheen, M., VanCompernolle, M., Arrowsmith, L. M., Peel, L. R., Queiroz, N., Vedor, M., da Costa, I., Mucientes, G., Couto, A., Humphries, N. E., Abalo-Morla, S., Abascal, F. J., Abercrombie, D. L., Abrantes, K., Abreu-Grobois, F. A.,… Eguíluz, V. M. (2025). Global tracking of marine megafauna space use reveals how to achieve conservation targets. Science, 388(6751), 1086–1097. 10.1126/science.adl0239

Södergren, K., & Palm, J. (2021). The role of local governments in overcoming barriers to industrial symbiosis. Cleaner Environmental Systems, 2, 100014. 10.1016/j.cesys.2021.100014

Stankovic, M., Mishra, A. K., Rahayu, Y. P., Lefcheck, J., Murdiyarso, D., Friess, D. A., Corkalo, M., Vukovic, T., Vanderklift, M. A., Farooq, S. H., Gaitan-Espitia, J. D., & Prathep, A. (2023). Blue carbon assessments of seagrass and mangrove ecosystems in south and Southeast Asia: Current progress and knowledge gaps. Science of The Total Environment, 904, 166618. 10.1016/j.scitotenv.2023.166618

Sunderland, T., Sunderland-Groves, J., Shanley, P., & Campbell, B. (2009). Bridging the Gap: How Can Information Access and Exchange Between Conservation Biologists and Field Practitioners be Improved for Better Conservation Outcomes? Biotropica, 41(5), 549–554. 10.1111/j.1744-7429.2009.00557.x

Sutherland, W. J., Pullin, A. S., Dolman, P. M., & Knight, T. M. (2004). The need for evidence-based conservation. Trends in Ecology & Evolution, 19(6), 305–308. 10.1016/j.tree.2004.03.018

Tanalgo, K. C. (2025). Open and FAIR data sharing are building blocks to bolster biodiversity conservation in Southeast Asia. Biological Conservation, 307, 111192. 10.1016/j.biocon.2025.111192

Tetley, M.J., Braulik, G.T., Lanfredi, C., Minton, G., Panigada, S., Politi, E., Zanardelli, M., Notarbartolo di Sciara, G. and Hoyt, E., 2022. The important marine mammal area network: a tool for systematic spatial planning in response to the marine mammal habitat conservation crisis. Frontiers in Marine Science, 9, p.841789.

UNEP-WCMC, 2026. State of the World’s Migratory Species: Interim Report (2026). UNEP-WCMC, Cambridge, United Kingdom.

Zenni, R. D., Barlow, J., Pettorelli, N., Stephens, P., Rader, R., Siqueira, T., Gordon, R., Pinfield, T., & Nuñez, M. A. (2023). Multi-lingual literature searches are needed to unveil global knowledge. Journal of Applied Ecology, 60(3), 380–383. 10.1111/1365-2664.14370

