## Supplemental Material for "What we miss shapes what we protect: sea turtles as a case study of grey literature and language bias in an evidence synthesis"

### Method Description A: Literature search details for green turtle across Southeast Asia

To systematically retrieve relevant literature, we developed English-language search strings based on previous search strategies used in marine megafauna studies (Kot et al., 2023a). The search strategy was designed to capture literature using the following categories of terms:

i. Focal species – taxonomic and common names of green turtles and dugongs
ii. Study type – sampling methodologies related to movement
iii. Evidence of connectivity – keywords related to behaviours, activities, sites associated with migration and movement
iv. Geographic region – names of relevant countries, territories, and marine areas

The initial search string was structured as follows: ("Chelonia mydas" OR “green turtle” OR "green sea turtle" OR dugong) AND (satellite OR telemetry OR tag* OR track* OR “photo ID” OR isotop* OR genetic* OR mark OR recapture) AND (migrat* OR connect* OR move* OR feed* OR forag* OR breed* OR dispers* OR nest* OR travel* OR aggregat* OR ground* OR corridor* OR route* OR habitat* OR distribut* OR barrier* OR releas* OR arriv*) AND (Indonesia OR Malaysia OR Cambodia OR Thailand OR Philippines OR Vietnam OR Taiwan OR China OR “Hong Kong” OR “Gulf of Thailand” OR “South China Sea” OR “West Philippine Sea” OR Asia* OR “Andaman Sea” OR “Strait of Malacca” OR “Malacca Strait”). The search string included both ‘green turtles’ and ‘dugongs’ as they are sympatric species. However, only two dugong-tracking studies met the criteria for meta-analysis, indicating that over the past 38 years, research on movement has mainly concentrated on green turtles, with limited attention to dugongs. This imbalance leads us to exclude dugong references from subsequent analyses, focusing exclusively on green turtles.

The initial search strings translated into each regional language were as follows:

1) Filipino: ("Chelonia mydas" OR "pawikan" OR “green turtle” OR dugong) AND (satellite OR telemetry OR tag OR sundan OR "photo ID" OR isotope OR genetic OR mark OR "muling makuha") AND (lumipat OR kumonekta OR gumalaw OR magpakain OR kumain OR lahi OR kumalat OR pugad OR paglalakbay OR pagsasama OR lupa OR corridor OR ruta OR tirahan OR ipamahagi OR hadlang OR pakawalan OR dumating) AND (Indonesia OR Malaysia OR Cambodia OR Thailand OR Pilipinas OR Vietnam OR Taiwan OR Tsina OR "Hong Kong" OR

"Gulf of Thailand" OR "South China Sea" OR "West Philippine Sea" OR Asya OR "Andaman Sea" OR "Strait ng Malacca")

2) Indonesian: ("Chelonia mydas" OR "penyu hijau" OR *dugong) AND (satelit OR telemetri OR menandai OR melacak OR "foto ID" OR isotop OR genetik OR tanda OR "menangkap kembali") AND (*migrasi OR *hubung OR *pindah OR "memberi makan" OR "mencari makan" OR "berkembangbiak'" OR *sebar OR *sarang OR *jalan OR agregat OR tanah OR rute OR habitat OR *distribusi OR *halang OR *lepas OR tiba) AND (Indonesia OR Malaysia OR Kamboja OR Thailand OR Filipina OR Vietnam OR Taiwan OR Cina OR "Hong Kong" OR "Teluk Thailand" OR "Laut Cina Selatan" OR "Laut Filipina Barat" OR Asia OR "Laut Andaman" OR "Selat Malaka")

3) Khmer: ("Chelonia mydas" OR ល្មិច OR “Dugong dugon” OR ជ្រូកទឹក) AND (ផ្កាយរណប OR ទូរមាត្រ OR ស្លាក OR តាមដាន OR "រូបសម្គាល់អត្តសញ្ញាណ" OR isotope OR ហ្សែន OR ស្នាម OR "ការចាប់យកឡើងវិញ") AND (បំលាស់ទី OR ទំនាក់ទំនង OR ផ្លាស់ប្តូរ OR ចំណី OR រកចំណី OR ពូជ OR បំបែក OR សំបុក OR ធ្វើដំណើរ OR ប្រមូលផ្តុំ OR ដី OR ច្រករបៀង OR ផ្លូវ OR ទីជម្រក OR ចែកចាយ OR របាំង OR ដោះលែង OR មកដល់) AND (ឥណ្ឌូនេស៊ី OR ម៉ាឡេស៊ី OR កម្ពុជា OR ថៃ OR ហ្វីលីពីន OR វៀនណាម OR តៃវ៉ាន់ OR ចិន OR ហុងកុង OR "ឈូងសមុទ្រថៃ" OR "សមុទ្រចិនខាងត្បូង" OR "សមុទ្រហ្វីលីពីនខាងលិច" OR អាស៊ី OR "សមុទ្រអាន់ដាម៉ាន់" OR "ច្រកសមុទ្រម៉ាឡាកា")

4) Malay: ("Chelonia mydas" OR "penyu agar" OR dugong) AND (satelit OR telemetri OR tag OR jejak OR "foto ID" OR isotop OR genetik OR tanda OR "tangkap semula") AND (*hijrah OR *hubung OR *gerak OR *makan OR "cari makan" OR *biak OR sebar OR *sarang OR perjalanan OR *kumpul OR tapak OR koridor OR laluan OR habitat OR *tabur OR rintangan OR *lepas OR tiba) AND (Indonesia OR Malaysia OR Kemboja OR Thailand OR Filipina OR Vietnam OR Taiwan OR China OR "Hong Kong" OR "Teluk Thailand" OR "Laut Cina Selatan" OR "Laut Filipina Barat" OR "Asia" OR "Laut Andaman” OR "Selat Melaka")

5) Thai: ("Chelonia mydas" OR เต่าตนุ OR “Dugong dugon” OR พะยูน) AND (ดาวเทียม OR มาตรวัดระยะไกล OR ป้าย OR ติดตาม OR รูปถ่ายระบุตัวตน OR isotope OR พันธุกรรม OR เครื่องหมาย OR จับซ้ำ) AND (*อพยพ OR *สัมพันธ์ OR *เคลื่อน OR *กิน OR *หาอาหาร OR *สืบพันธุ์ OR แพร่ OR *ที่อยู่ OR *เดินทาง OR *รวบรวม OR บริเวณ OR *แนวเชื่อมต่อ OR เส้นทาง OR *ที่อยุ่อาศัย OR *กระจาย OR *กีดขวาง OR *ปล่อย OR *ถึง) AND (อินโดนีเซีย OR มาเลเซีย OR กัมพูชา OR ไทย OR ฟิลิปปินส์ OR เวียดนาม OR ไต้หวัน OR จีน OR ฮ่องกง OR อ่าวไทย OR ทะเลจีนใต้ OR ทะเลฟิลิปปินส์ตะวันตก OR เอเชีย OR ทะเลอันดามัน OR ช่องแคบมะละกา)

6) Traditional Chinese: ("Chelonia mydas" OR 綠蠵龜 OR “Dugong dugon” OR 儒艮) AND (衛星 OR 遙測 OR 標 OR 追蹤 OR 影像辨識 OR 同位素 OR 基因 OR 標記 OR 再補捉) AND (遷移 OR 連結 OR 移動 OR 覓食 OR 繁殖 OR 散佈 OR 卵窩 OR 旅行 OR 聚集 OR 地 OR 走廊 OR 路線 OR 棲息地 OR 分佈 OR 屏障 OR 野放 OR 抵達) AND (印尼 OR 馬來西亞 OR 柬埔寨 OR 泰國 OR 菲律賓 OR 越南 OR 台灣 OR 中國 OR 香港 OR 泰國灣 OR 南中國海 OR 西菲律賓海 OR 亞洲 OR 安達曼海 OR 馬六甲海峽)
7) Vietnamese: ("Chelonia mydas" OR "Rùa xanh" OR “Dugong dugon” OR "Bò Biển") AND ("Vệ tinh" OR "Dữ liệu từ xa" OR "Nhãn" OR "Tuyến" OR "Nhận diện hình ảnh" OR isotope OR "di chuyển" OR "đánh đấu" OR "đánh bắt lại") AND ("di cư" OR "kết đôi" OR "di chuyển" OR "kiếm ăn" OR "hành vi kiếm ăn" OR "sinh sản" OR "sự phân tán" OR "tổ sinh sản" OR "Chuyến đi" OR "tập chung" OR "bãi rùa đẻ" OR "hành lang" OR "hành trình di cư" OR "sinh cảnh" OR "Phân bố" OR "rào cản" OR "tái thả" OR "định cư") AND (Indonesia OR Malaysia OR Campuchia OR "Thái Lan" OR Philippines OR "Việt Nam" OR "Đài Loan" OR "Trung Quốc" OR "Hong Kong" OR "Vịnh Thái Lan" OR "Biển Đông" OR "Đông Nam Á" OR "Châu Á" OR "Biển Andaman" OR "Eo biển Malacca")

The results of the literature search were imported into Covidence (www.covidence.org), a software designed to streamline the systematic review process. Each reference was first screened based on its title and abstract and assigned as “Relevant”, “Not Relevant” or “Maybe Relevant” according to the following inclusion criteria: (i) the paper contains telemetry (satellite/acoustic) data, (ii) the paper identifies two or more locations indicating movement or migration, (iii) the paper describes repeated sightings of an individual across time and space, or (iv) the paper identifies a ‘stopover site’ (a site en route to the primary foraging ground, presumably for feeding or resting; Perez et al., 2022) in migratory routes. References assigned as “Relevant” or “Maybe Relevant” were sent for full-text review for confirmation. At this stage, references marked as “Maybe Relevant” were independently reviewed by another reviewer for a second assessment. If at least one of the two reviewers deemed the study relevant, it was included for further analysis.

**Method Description B: Creation of regional and sub-regional connectivity models**

Individual georeferenced sites extracted from different studies of the same species were aggregated into ‘metasites’ through a three-step process: data preparation, site aggregation, and metasite creation. We iteratively selected and grouped individual sites by creating buffers around the centroids following a set of rules, including 1) maintaining high resolutions for reproductive sites with a 15 km radius buffer and applying larger radii (50 km for national-scale and 100 km for regional-scale) to non-reproductive sites,

2) differentiating reproductive sites from non-reproductive sites by separating them in the aggregation process, and 3) retaining information on large-scale (>500 km) connections resulting from animal movements. These rules were applied to ensure that the resolution of information on both critical reproductive life history stages and large-scale movements was not lost during aggregation.

After sorting the sites in the dataset by connection distance, a site (the 'Lookup Site') was selected iteratively to determine which other sites could be aggregated with it. These were sites ('Nearby Sites') that fell within the Lookup Site's buffer. Any Nearby Site with a route or connection >500 km Euclidean distance was removed from grouping with the Lookup Site (now 'Removed Site'). The Removed Site was then grouped with its own Nearby Site(s) down the pipeline. Any successful groupings formed a metasite, with a new centroid calculated as the average latitude and longitude of the Lookup Site and Nearby Site(s), and the Lookup Site's buffer size remaining as the metasite buffer. New metasites were created repeatedly until all sites intersected the metasite buffer and were aggregated into respective metasites. Subsequently, all connections linking the same metasites were joined into metaconnections. To avoid underestimating the contribution of non-English and grey literature to the understanding of migratory connectivity, we included all sites without removing potential duplicates across references to ensure that each connectivity information was preserved, similar to Kot et al. (2023a). This approach accounts for the discrepancies in transparency and detail among scientific articles and grey literature, especially regarding mark-recapture data, which often lack individual or geographic details. Consequently, the strength of connectivity presented in this study, which was calculated based on the number of individuals using a pathway, reflects reported individuals rather than unique individuals per se. Final regional model outputs comprised points (metasites) and polylines (metaconnections) showing directions of migratory movements between Exclusive Economic Zones (EEZs), while sub-regional models show migratory links between distinct sites within EEZs.

**Table S1** List of regional publication databases:

| No | Database Name | Region | URL |
|---|---|---|---|
| 1 | Asean Citation Index | ASEAN | https://asean-cites.org/ |
| 2 | Malaysian Citation Index (MYCITE) | Malaysia | https://mycite.mohe.gov.my/ |
| 3 | MyJurnal | Malaysia | https://myjurnal.mohe.gov.my/ |
| 4 | Airiti Library | Taiwan | https://www.airitilibrary.com/ |
| 5 | National Digital Library of Theses and Dissertations in Taiwan | Taiwan | https://ndltd.ncl.edu.tw/ |
| 6 | Sinta (Science and Technology Index) | Indonesia | https://sinta.kemdikbud.go.id/ |
| 7 | Perpustakaan Nasional | Indonesia | https://edeposit.perpusnas.go.id/collection |
| 8 | Garba Rujukan Digital (GARUDA) | Indonesia | https://garuda.kemdikbud.go.id/ |
| 9 | Vietnam Science and Technology Publication Database (VISTA) | Vietnam | https://sti.vista.gov.vn/Pages/Trang-chu.aspx |
| 10 | VAST | Vietnam | http://elib.isivast.org.vn/ |
| 11 | LIC-VNU | Vietnam | https://lic.vnu.edu.vn/vi |
| 12 | PMBC Research Bulletin | Thailand | https://km.dmcr.go.th/c_266 |
| 13 | ThaiLIS Digital Collection (TDC) | Thailand | https://tdc.thailis.or.th/ |
| 14 | OpenDevelopment Mekong | Mekong Region (Cambodia, Laos, Thailand, Vietnam, and Myanmar) | https://opendevelopmentmekong.net/background/ |
| 15 | Cambodian Journal for Natural History | Cambodia | http://www.rupp.edu.kh/cjnh/index.php |
| 16 | Philippine E-Journals | Philippines | https://ejournals.ph/index.php |
| 17 | Transactions of the National Academy of Science and Technology Philippines (Journal) | Philippines | https://transactions.nast.ph/ |
| 18 | Science Diliman (Journal) | Philippines | https://journals.upd.edu.ph/index.php/sciencediliman |
| 19 | UPLB Journals Online | Philippines | https://ovcre.uplb.edu.ph/journals-uplb/ |

**Table S2** List of professional organisations and databases. This list is non-exhaustive and does not include all professional contacts consulted during the literature search.

| No | Organisation Name | Region | Resource Type |
|---|---|---|---|
| 1 | IUCN-SSC Marine Turtle Specialist Group | Global | Regional reports |
| 2 | Global Biodiversity Information Facility (GBIF) | Global | Biodiversity data portal |
| 3 | Sea Turtle Document Library (hosted on seaturtle.org) | Global | Journal articles, conference proceedings, theses, technical reports |
| 4 | Ocean Biogeographic Information System Spatial Ecological Analysis of Megavertebrate Population (OBIS-SEAMAP) | Global | Data portal |
| 5 | Migratory Connectivity in the Ocean (MiCO) system | Global | Data portal |
| 6 | Marine Turtle Breeding and Migration Atlas (TurtleNet) | Global | Data portal |
| 7 | Movebank | Global | Data portal |
| 8 | Southeast Asian Fisheries Development Center (SEAFDEC) | Regional | Technical reports, conference proceedings, journal articles |
| 9 | Joint Management Committee of the Turtle Islands Heritage Protected Area (TIHPA) | Regional | Unpublished data |
| 10 | Fisheries Research Institute, Department of Fisheries Malaysia | Malaysia | Technical reports, conference proceedings, journal articles |
| 11 | State government agencies (Sabah Park, Sabah Wildlife Department, Sabah Biodiversity Centre, Sarawak Forestry Corporation) | Malaysia | Annual reports, strategic plans, unpublished data |
| 12 | Malaysia Biodiversity Information System (MyBIS) | Malaysia | Biodiversity data portal |
| 13 | NGOs (WWF-Malaysia, Marine Research Foundation, Lang Tengah Turtle Watch, etc.) | Malaysia | Annual reports, newsletters, technical reports, unpublished data |
| 14 | Sea Turtle Research Unit (SEATRU), Universiti Malaysia Terengganu | Malaysia | Unpublished data |
| 15 | County and city governments (Ocean Conservation Administration) | Taiwan | Technical reports |
| 16 | Taiwan Biodiversity Information Alliance (TBIA) Data Portal | Taiwan | Biodiversity data portal |
| 17 | TurtleSpot Taiwan | Taiwan | Unpublished data |
| 18 | Ministry of Marine Affairs and Fisheries (Kementerian Kelautan dan Perikanan Republik Indonesia, KKP) | Indonesia | Annual reports, books |
| 19 | National Research and Innovation Agency (Badan Riset dan Inovasi Nasional, BRIN) | Indonesia | Annual reports, journal articles |
| 20 | NGOs (WWF-Indonesia, Yayasan Pulau Banyak, Thrive Conservation, Anambas Foundation, etc.) | Indonesia | Annual reports, journal articles, unpublished data |
| 21 | The Marine and Coastal Resources Research Center, Department of Marine and Coastal Resources (DMCR) | Thailand | Technical reports |
| 22 | Department of Fisheries | Thailand | Technical reports |
| 23 | Research Institute for Marine Fisheries | Vietnam | Magazines, technical reports, books |

| | | | |
|---|---|---|---|
| 24 | National Bureau of Science and Technology Information | Vietnam | Technical reports, journal articles |
| 25 | Institute of Ecology and Biological Resources | Vietnam | Conference proceedings |
| 26 | Vietnam Academy of Science & Technology | Vietnam | Annual reports, newsletters |
| 27 | Centre for Biodiversity and Endangered Species | Vietnam | Annual reports |
| 28 | Institute of Oceanography Nha Trang | Vietnam | Book, technical reports, conference proceedings, journal articles |
| 29 | Khmer Ocean Life | Cambodia | Unpublished data |
| 30 | OpenDevelopment Mekong | Cambodia | Technical reports, action plans |
| 31 | Fauna & Flora International Cambodia | Cambodia | Technical reports, unpublished data |
| 32 | Agriculture, Fisheries and Conservation Department (AFCD), Hong Kong | Hong Kong | Annual reports, unpublished data |
| 33 | Department of Environment and Natural Resources, Biodiversity Management Bureau, Philippines | Philippines | Technical reports, unpublished data |
| 34 | Large Marine Vertebrates Research Institute Philippines (LAMAVE) | Philippines | Unpublished data |

123

**Table S3** The number of references with connectivity studies conducted in each jurisdiction (number of references published by that jurisdiction in parentheses).

| Jurisdiction | Journal Articles | Newsletters | Dissertations | Reports | Proceedings | Books | Online Databases | Unpublished Data | Total References | Reference Types |
|---|---|---|---|---|---|---|---|---|---|---|
| Malaysia | 8 (8) | 4 (4) | 2 (1) | 10 (2) | 10 (1) | 0 (0) | 6 (5) | 4 (3) | 44 (21) | 7 (7) |
| Taiwan | 8 (8) | 0 (0) | 5 (5) | 4 (4) | 3 (0) | 0 (0) | 0 (0) | 4 (3) | 24 (17) | 5 (4) |
| Thailand | 3 (3) | 0 (0) | 1 (0) | 5 (0) | 10 (1) | 1 (0) | 0 (0) | 1 (0) | 20 (4) | 5 (2) |
| Indonesia | 2 (2) | 2 (2) | 1 (1) | 2 (0) | 0 (0) | 1 (1) | 8 (8) | 2 (0) | 18 (14) | 7 (5) |
| Philippines | 3 (3) | 3 (3) | 0 (0) | 4 (1) | 3 (1) | 0 (0) | 2 (1) | 3 (0) | 18 (9) | 6 (5) |
| Vietnam | 2 (2) | 0 (0) | 1 (1) | 4 (1) | 1 (0) | 1 (0) | 0 (0) | 1 (1) | 10 (4) | 6 (4) |
| Hong Kong | 2 (2) | 1 (1) | 0 (0) | 0 (0) | 0 (0) | 2 (1) | 0 (0) | 1 (2) | 6 (4) | 4 (4) |
| Cambodia | 0 (0) | 0 (0) | 0 (0) | 3 (0) | 0 (0) | 0 (0) | 0 (0) | 0 (0) | 3 (0) | 1 (0) |
| Federated States of Micronesia | 2 (2) | 1 (1) | 0 (0) | 2 (0) | 1 (0) | 0 (0) | 0 (0) | 1 (NA) | 6 (3) | 4 (2) |
| China | 2 (2) | 2 (2) | 0 (0) | 1 (0) | 0 (0) | 0 (0) | 0 (0) | 0 (NA) | 5 (4) | 3 (2) |
| Japan | 1 (1) | 0 (0) | 0 (1) | 1 (0) | 2 (15) | 0 (0) | 0 (0) | 1 (NA) | 5 (17) | 4 (3) |
| Guam | 0 (0) | 0 (0) | 0 (0) | 2 (0) | 2 (0) | 0 (0) | 0 (0) | 0 (NA) | 4 (0) | 2 (0) |
| Republic of Palau | 0 (0) | 1 (1) | 0 (0) | 2 (0) | 1 (0) | 0 (0) | 0 (0) | 0 (NA) | 4 (1) | 3 (1) |
| Republic of the Marshall Islands | 1 (1) | 0 (0) | 0 (0) | 2 (0) | 0 (0) | 0 (0) | 0 (0) | 0 (NA) | 3 (1) | 2 (1) |
| Myanmar | 0 (0) | 0 (0) | 0 (0) | 1 (0) | 0 (0) | 0 (0) | 0 (0) | 0 (NA) | 1 (0) | 1 (0) |
| Australia | 1 (1) | 0 (0) | 0 (1) | 0 (0) | 0 (0) | 0 (0) | 0 (0) | 0 (NA) | 1 (2) | 1 (2) |
| Papua New Guinea | 0 (0) | 0 (0) | 0 (0) | 1 (0) | 0 (0) | 0 (0) | 0 (0) | 0 (NA) | 1 (0) | 1 (0) |

| Jurisdiction | Journal Articles | Newsletters | Dissertations | Reports | Proceedings | Books | Online Databases | Unpublished Data | Total References | Reference Types |
|---|---|---|---|---|---|---|---|---|---|---|
| New Caledonia | 0 (0) | 0 (0) | 0 (0) | 0 (0) | 1 (0) | 0 (0) | 0 (0) | 0 (NA) | 1 (0) | 1 (0) |
| Italy | 0 (0) | 0 (0) | 0 (0) | 0 (0) | 0 (1) | 0 (0) | 0 (0) | 0 (NA) | 0 (1) | 0 (1) |
| Switzerland | 0 (0) | 0 (0) | 0 (0) | 0 (0) | 0 (0) | 0 (1) | 0 (0) | 0 (NA) | 0 (1) | 0 (1) |
| United States of America | 0 (0) | 0 (0) | 0 (0) | 0 (1) | 0 (3) | 0 (0) | 0 (0) | 0 (NA) | 0 (4) | 0 (2) |
| Intergovernmental organisation | 0 (0) | 0 (0) | 0 (0) | 0 (8) | 0 (0) | 0 (0) | 0 (0) | 0 (NA) | 0 (8) | 0 (1) |

**Table S4** The number of connected territories for each focal jurisdiction, and the loss of connected jurisdiction when considering only peer-reviewed, English-language literature. The overall network impact is calculated by multiplying the number of lost jurisdictions by the number of connected jurisdictions. Since Hong Kong does not have a separate EEZ, it is reported under China.

| Focal Jurisdiction | Number of Connected Jurisdictions | Connected Jurisdictions | Number of Lost Jurisdictions | Lost Jurisdictions | % of Paired Jurisdictions Lost | % of Total Network Pairs Lost | Overall Network Impact |
|---|---|---|---|---|---|---|---|
| *All literature vs Peer-reviewed-literature only* | | | | | | | |
| Philippines | 14 | China, Guam, Indonesia, Japan, Malaysia, Micronesia, New Caledonia, Northern Mariana Islands, Palau, Solomon Islands, Spratly Islands, Taiwan, Thailand, Vietnam | 6 | Guam, Japan, New Caledonia, Palau, Solomon Islands, Thailand | 42.9 | 11.5 | 84 |
| Thailand | 8 | Brunei, Cambodia, India, Indonesia, Malaysia, Philippines, Singapore, Vietnam | 6 | Brunei, Cambodia, Indonesia, Malaysia, Philippines, Singapore | 75 | 11.5 | 48 |
| Malaysia | 10 | Brunei, Indonesia, Micronesia, Palau, Papua New Guinea, Philippines, Singapore, Spratly Islands, Thailand, Vietnam | 4 | Palau, Papua New Guinea, Singapore, Thailand | 40 | 7.7 | 40 |
| Indonesia | 9 | Australia, Brunei, China, Malaysia, Palau, Philippines, Solomon Islands, Thailand, Vietnam | 4 | Brunei, Solomon Islands, Thailand, Vietnam | 44.4 | 7.7 | 36 |
| Vietnam | 8 | Cambodia, China, Indonesia, Malaysia, Philippines, Spratly Islands, Taiwan, Thailand | 2 | Indonesia, Spratly Islands | 25 | 3.8 | 16 |
| Taiwan | 5 | China, Japan, Micronesia, Philippines, Vietnam | 1 | Micronesia | 20 | 1.9 | 5 |
| Cambodia | 2 | Thailand, Vietnam | 1 | Thailand | 50 | 1.9 | 2 |
| China | 4 | Indonesia, Philippines, Taiwan, Vietnam | 0 | None | 0 | 0 | 0 |

| | | | | | | | |
|---|---|---|---|---|---|---|---|
| *All literature vs English-language-literature only* | | | | | | | |
| Taiwan | 5 | China, Japan, Micronesia, Philippines, Vietnam | 1 | Micronesia | 20 | 1.9 | 5 |
| Philippines | 14 | China, Guam, Indonesia, Japan, Malaysia, Micronesia, New Caledonia, Northern Mariana Islands, Palau, Solomon Islands, Spratly Islands, Taiwan, Thailand, Vietnam | 0 | None | 0 | 0 | 0 |
| Malaysia | 10 | Brunei, Indonesia, Micronesia, Palau, Papua New Guinea, Philippines, Singapore, Spratly Islands, Thailand, Vietnam | 0 | None | 0 | 0 | 0 |
| Indonesia | 9 | Australia, Brunei, China, Malaysia, Palau, Philippines, Solomon Islands, Thailand, Vietnam | 0 | None | 0 | 0 | 0 |
| Thailand | 8 | Brunei, Cambodia, India, Indonesia, Malaysia, Philippines, Singapore, Vietnam | 0 | None | 0 | 0 | 0 |
| Vietnam | 8 | Cambodia, China, Indonesia, Malaysia, Philippines, Spratly Islands, Taiwan, Thailand | 0 | None | 0 | 0 | 0 |
| China | 4 | Indonesia, Philippines, Taiwan, Vietnam | 0 | None | 0 | 0 | 0 |
| Cambodia | 2 | Thailand, Vietnam | 0 | None | 0 | 0 | 0 |

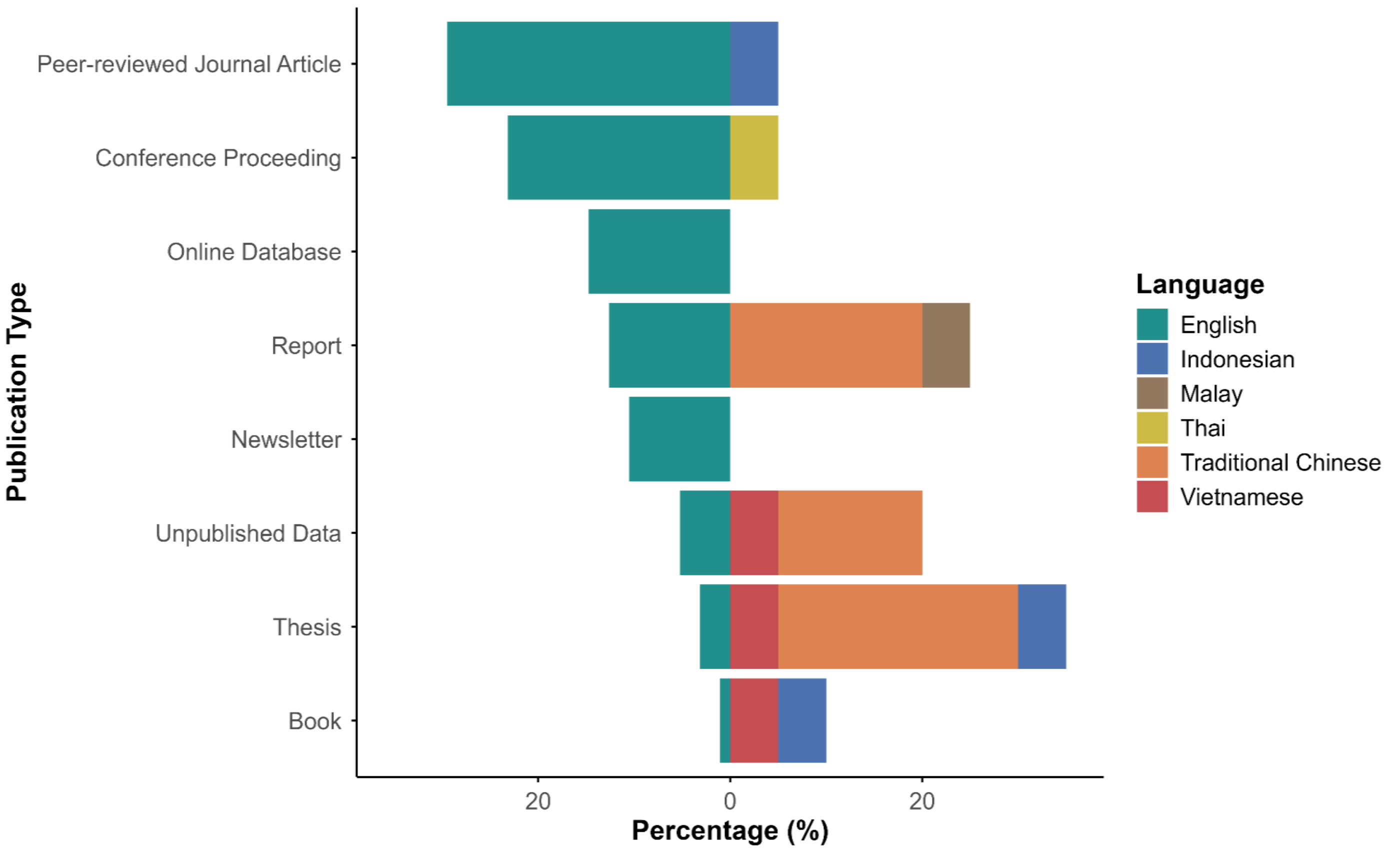


**Figure S1** Distribution of literature across English and non-English references.

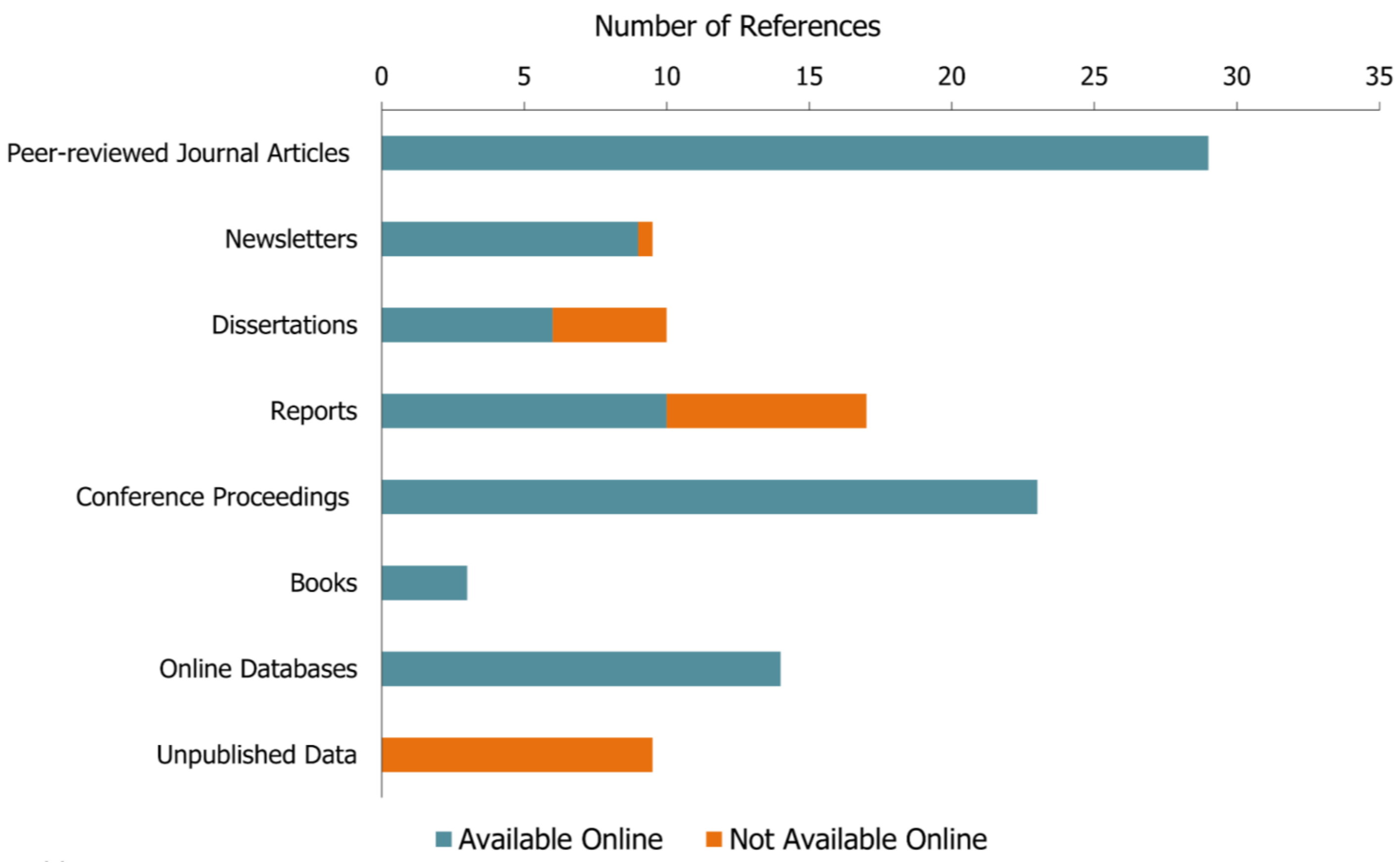


**Figure S2** Distribution of literature across the proportion of references available online.

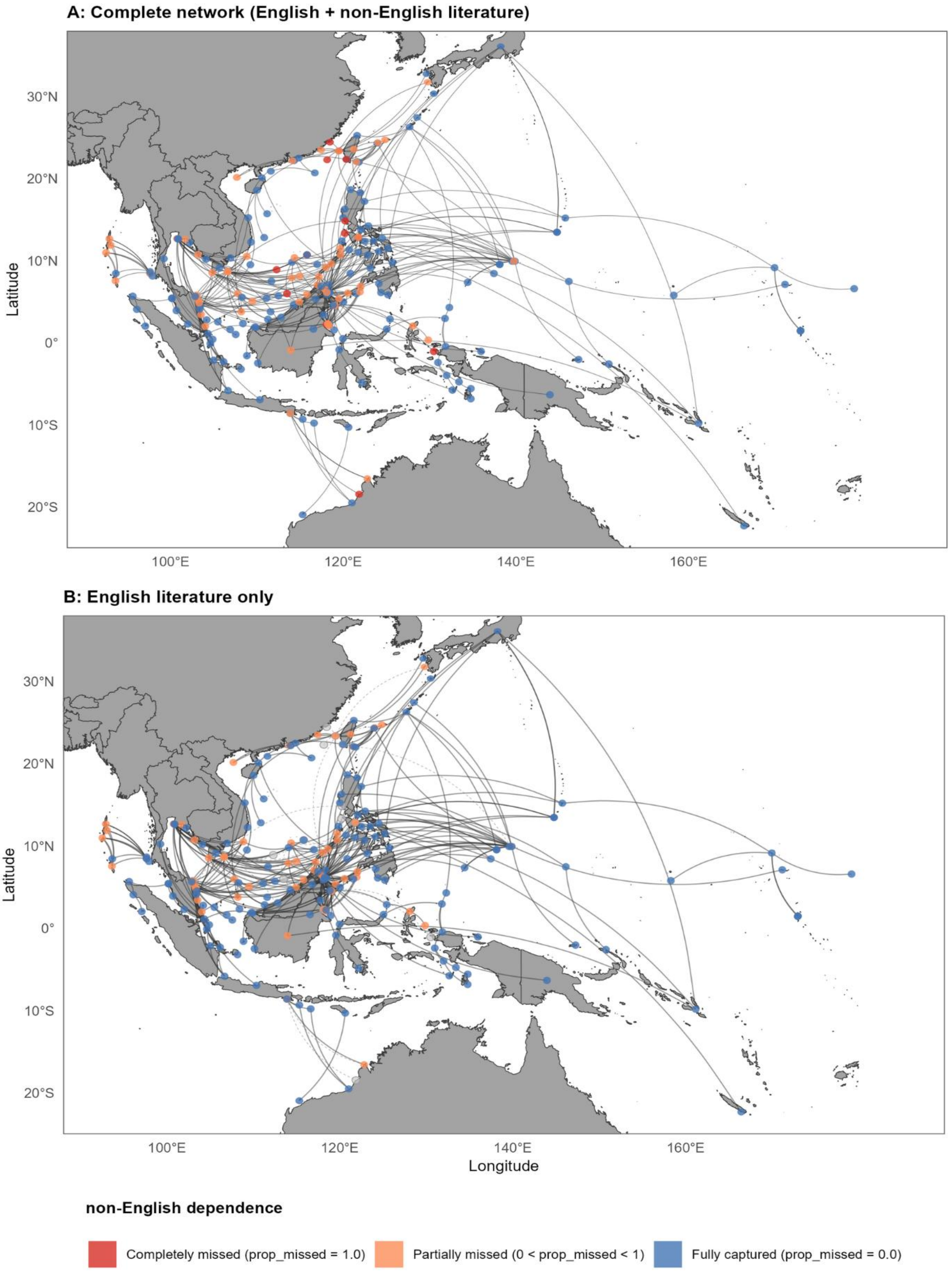
A: Complete network (English + non-English literature)
B: English literature only
30°N
20°N
10°N
0°
10°S
20°S
100°E
120°E
140°E
160°E
Latitude
Longitude
non-English dependence
Completely missed (prop_missed = 1.0)
Partially missed (0 < prop_missed < 1)
Fully captured (prop_missed = 0.0)

**Figure S3** Regional connectivity network of *Chelonia mydas* illustrating (A) the complete network with all literature included, and (B) network using only English literature. Circles represent metasites and the lines represent metaconnections. The visible network is colour-coded based on grey literature dependency. Missed metasites and metaconnections resulting from grey literature exclusion are shown in grey.

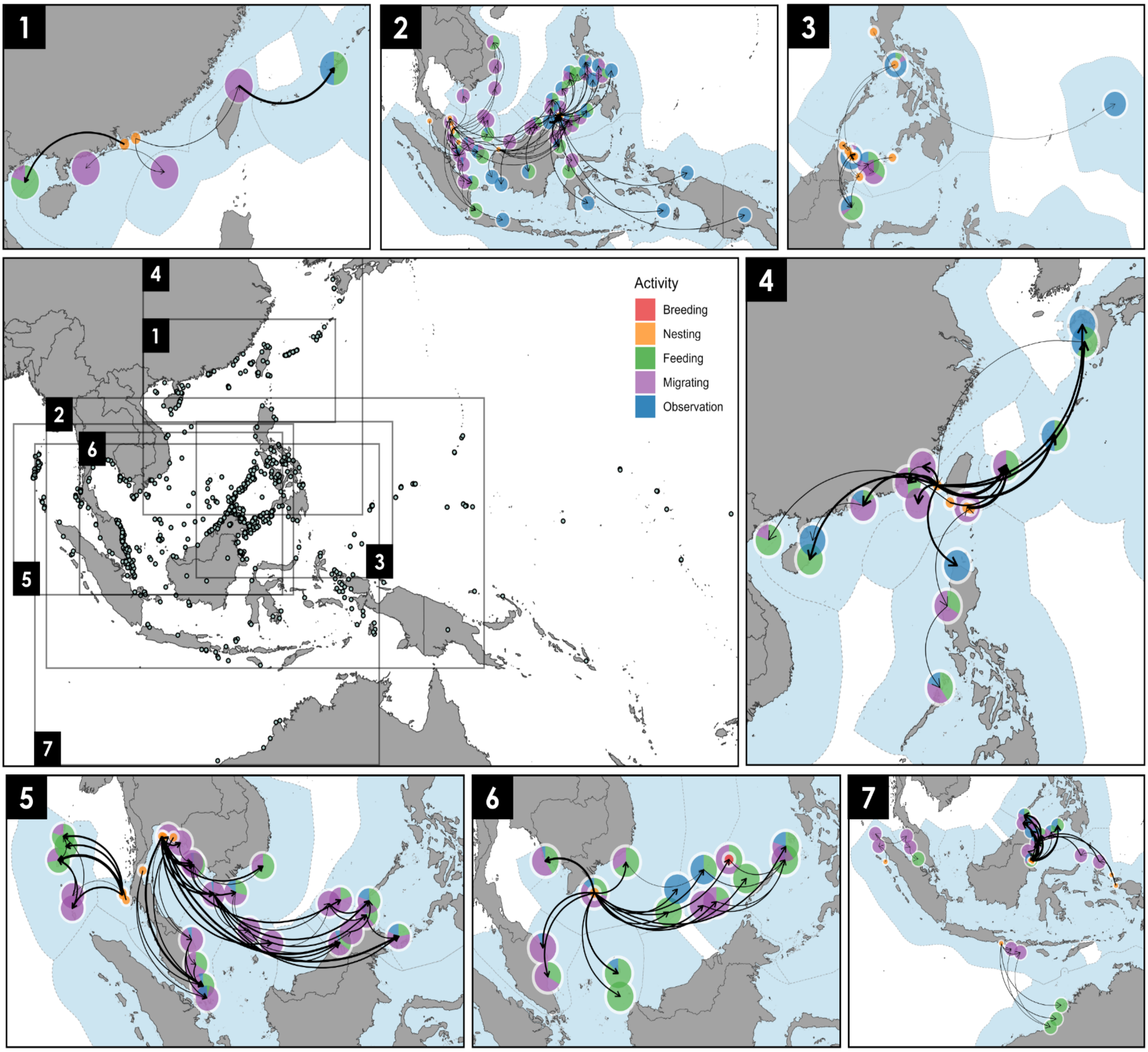


**Figure S4** Transboundary connectivity of *Chelonia mydas* **originating** from: (1) Hong Kong, (2) Malaysia, (3) the Philippines, (4) Taiwan, (5) Thailand, (6) Vietnam, and (7) Indonesia. No transboundary information is available for individuals tracked from Cambodia. The main panel shows the locations of all sites (circles) with connections. Nodes represent the animal's marked and recaptured locations, and the pie chart on each node indicates the proportion of activities (i.e., breeding, nesting, feeding, migrating, and observation). Node size reflects site type, with a smaller radius (15 km) for reproductive sites (breeding and nesting) and a larger radius (100 km) for non-reproductive sites (feeding, migrating and observation). Curved arrows represent connections, and line thickness indicates the scaled number of connections. Polygons around jurisdictions represent Exclusive Economic Zones (EEZs).

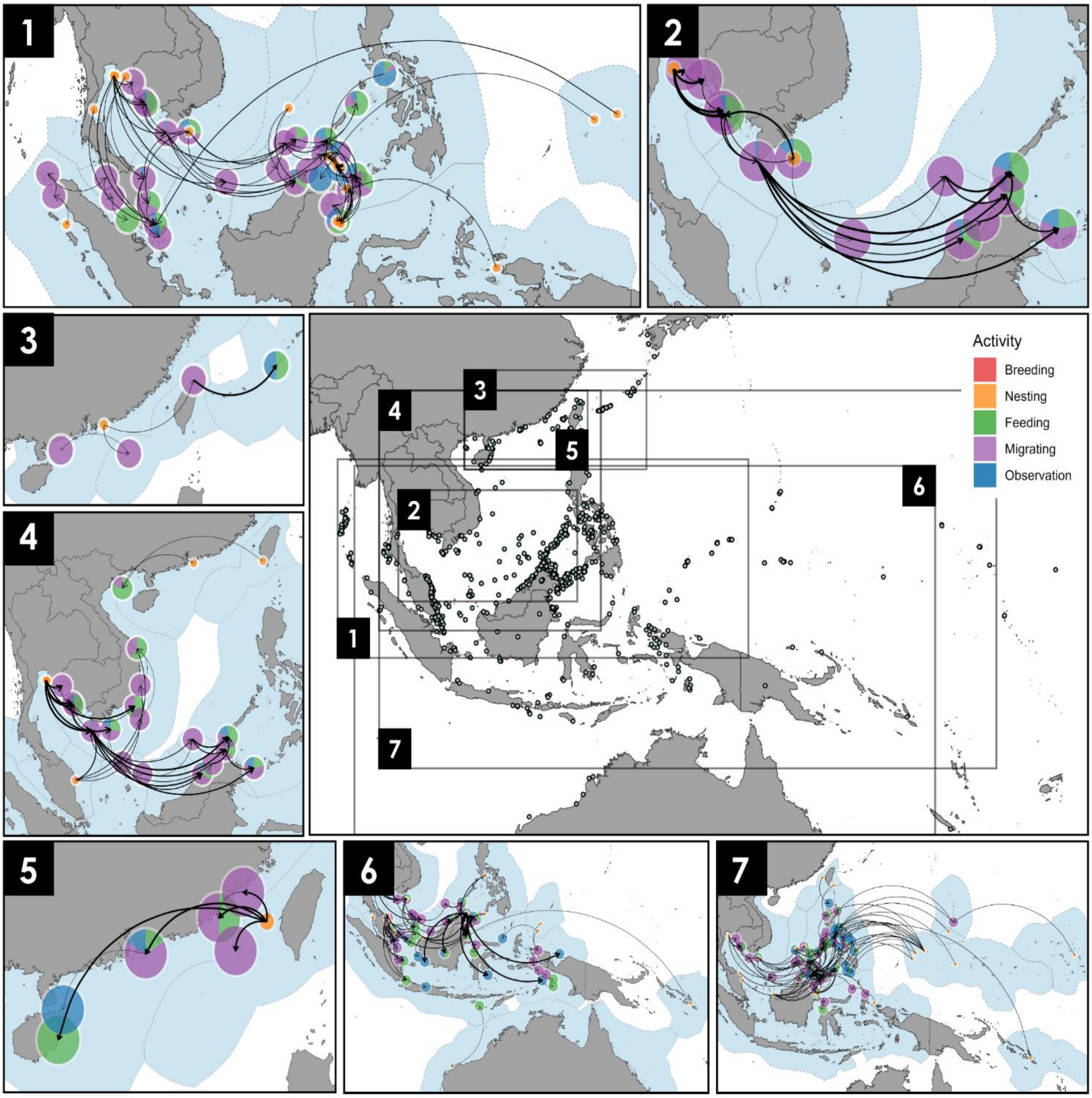


**Figure S5** Transboundary connectivity of *Chelonia mydas* **recaptured** in: (1) Malaysia, (2) Cambodia, (3) Taiwan, (4) Vietnam, (5) China, (6) Indonesia, and (7) the Philippines. No transboundary information is available for individuals recaptured in Thailand. The main panel shows the locations of all sites (circles) with connections. Nodes represent the animal's marked and recaptured locations, and the pie chart on each node indicates the proportion of activities (i.e., breeding, nesting, feeding, migrating, and observation). Node size reflects site type, with a smaller radius (15 km) for reproductive sites (breeding and nesting) and a larger radius (100 km) for non-reproductive sites (feeding, migrating and observation). Curved arrows represent connections, and line thickness indicates the scaled number of connections. Polygons around jurisdictions represent Exclusive Economic Zones (EEZs). Since Hong Kong does not have a separate EEZ, it is reported under China.

167

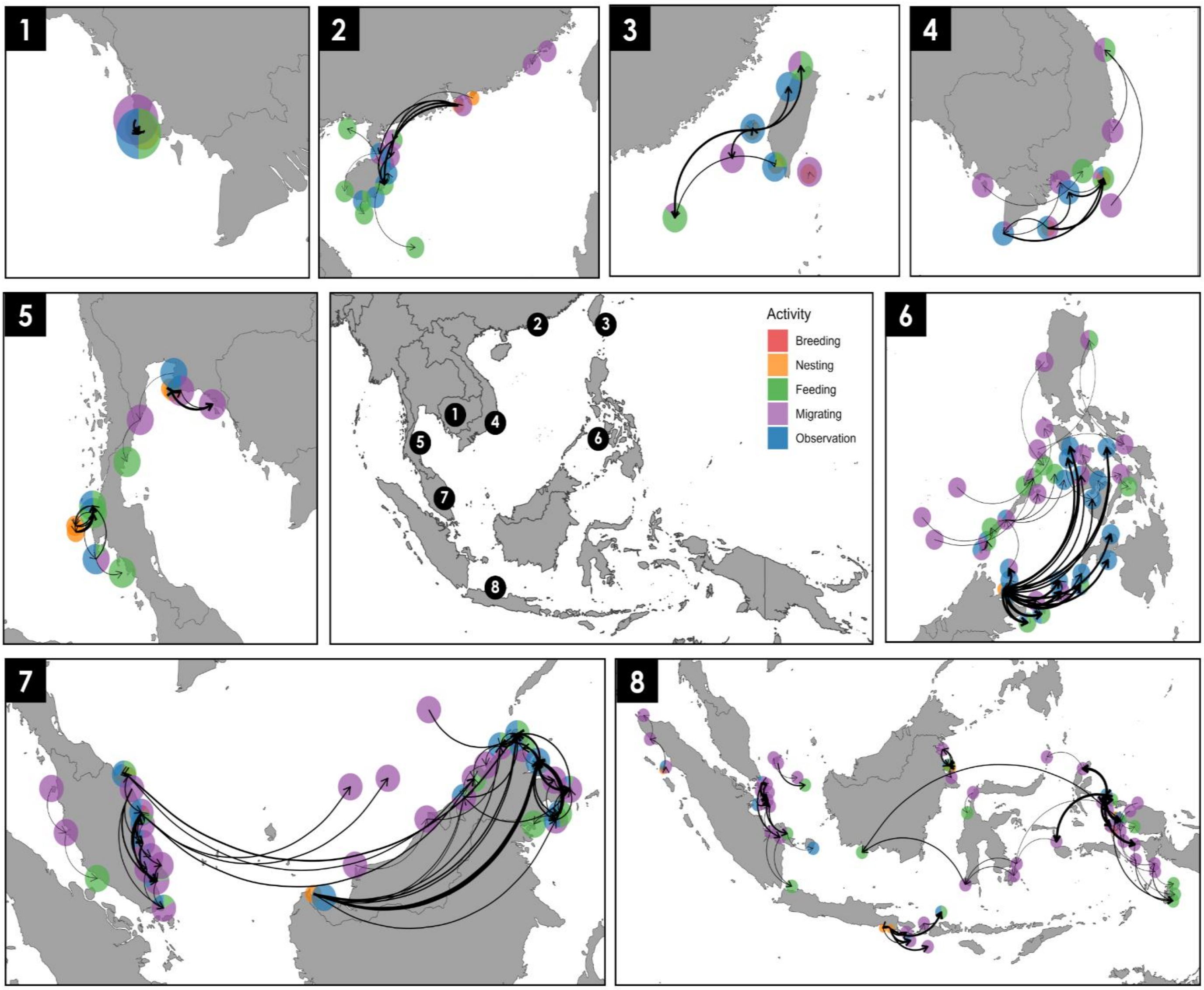


**Figure S6** In-jurisdiction connectivity of *Chelonia mydas* within: (1) Cambodia, (2) China, (3) Taiwan, (4) Vietnam, (5) Thailand, (6) the Philippines, (7) Malaysia, and (8) Indonesia. Nodes represent the animal's marked and recaptured locations, and the pie chart on each node indicates the proportion of activities (i.e., breeding, nesting, feeding, migrating, and observation). Node size reflects site type, with a smaller radius (15 km) for reproductive sites (breeding and nesting) and a larger radius (50 km) for non-reproductive sites (feeding, migrating and observation). Curved arrows represent connections, and line thickness indicates the scaled number of connections. Since Hong Kong does not have a separate EEZ, it is reported under China.

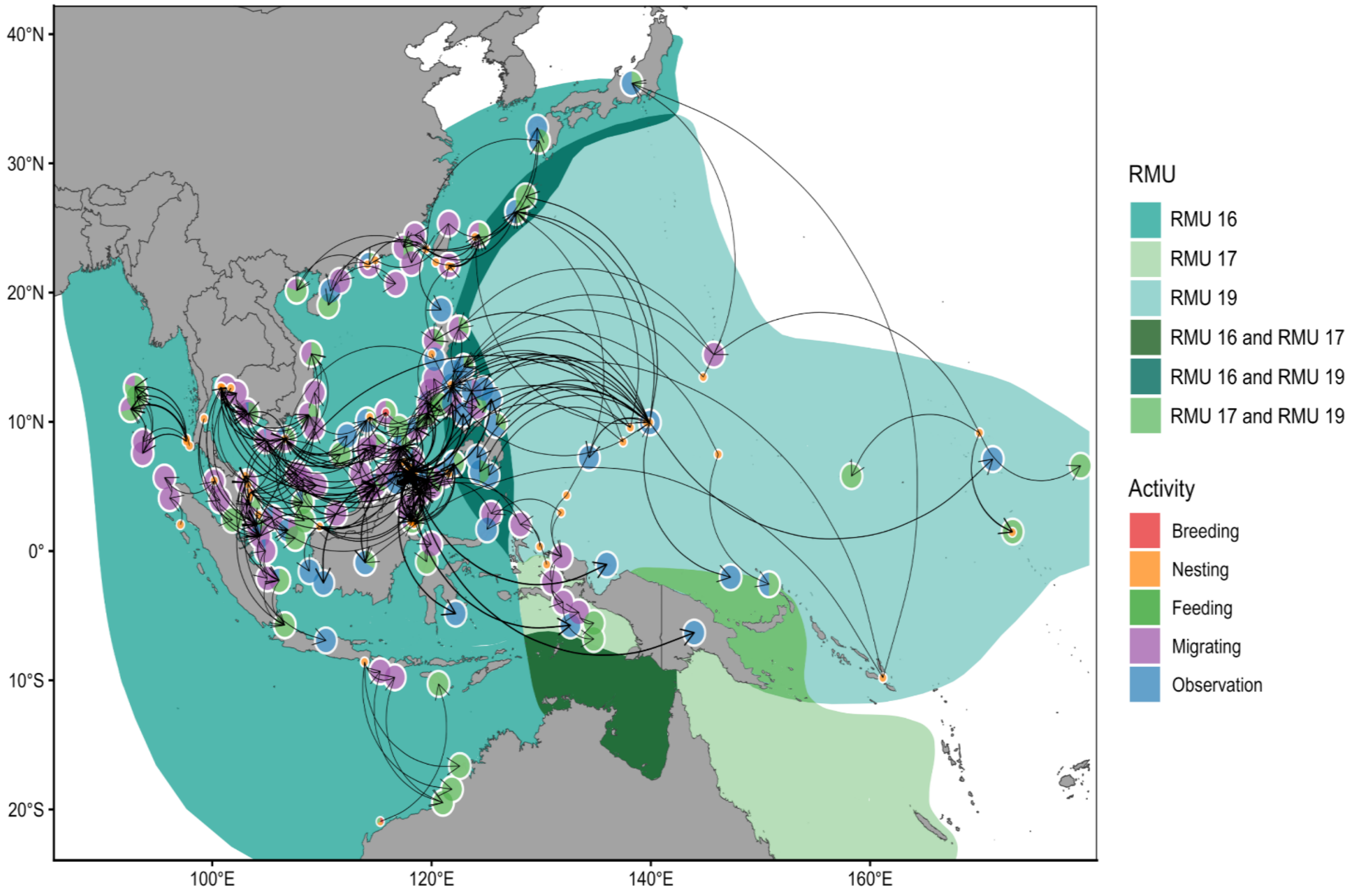


**Figure S7** Transboundary connectivity of *Chelonia mydas* illustrating migrations, with nesting/breeding connected to foraging and non-foraging sites across green turtle Regional Management Units (RMUs). RMU 16: East Indian and Southeast Asia; RMU 17: Southwest Pacific; RMU 19: West Central Pacific. Nodes, indicated by pie charts, represent metasites aggregated from individual sites, with segments showing the proportion of activity types at each metasite (i.e., breeding, nesting, feeding, migrating, and observation). Node size reflects site type, with a smaller radius (15 km) for reproductive sites (breeding and nesting) and a larger radius (100 km) for non-reproductive sites (feeding, migrating and observation). 'Observation' refers to those listed without a specifically documented behaviour or activity. Nodes on land are assigned to the EEZs of their respective territories without precise locations. Curved arrows indicate the movement directions, and line thickness reflects the scaled number of connections.
